# The pan-genome and local adaptation of *Arabidopsis thaliana*

**DOI:** 10.1101/2022.12.18.520013

**Authors:** Minghui Kang, Haolin Wu, Wenyu Liu, Mingjia Zhu, Yu Han, Wei Liu, Chunlin Chen, Kangqun Yin, Yusen Zhao, Zhen Yan, Huanhuan Liu, Shangling Lou, Yanjun Zan, Jianquan Liu

## Abstract

*Arabidopsis thaliana* has been used as a model species for research in a diverse collection of plant species. However, previous studies based on single reference genomes and short-read sequencing data are restricted to detecting variable genes and large structural variation (SV) underlying local adaptation. Here we *de novo* assemble high-quality chromosomal genomes of 38 *A. thaliana* ecotypes (with 6 relict ones) using PacBio-HiFi long-read sequencing. From these newly assembled genomes, we annotate several thousand new genes through pan-genomic analysis in comparison to the previous reference genome. The identified variable genes are mainly enriched in and associated with ecological adaptation and this species substantially expands its gene repertoire for local adaptation. We construct a graph-based pan-genome and identify 62,525 SVs which overlap with 14,243 genes. These genes are enriched in multiple ecological adaptation functions, including secondary metabolic processes, enzyme regulation, and biotic/abiotic stimulus. For example, a 566 bp insertion in the promoter of the light-adaptation *KNAT3* gene was specific to the high-altitude relict Tibet-0 ecotype. This SV reduces the expression level of *KNAT3* and promotes *A. thaliana* adaptation to habitats high in light radiation. In addition, compared with the SNPs, the SVs identified in this study captured the missing heritability and we detected novel SV associations with environmental variables in their native range, highlighting the value of SVs in environmental adaptation. The genome resources presented here will help pinpoint genetic changes that include both SVs and the ecotype-specific genes for local adaptation of *A. thaliana* and increase our understanding of the molecular mechanisms in this model species to respond to varied habitats.

## Introduction

*Arabidopsis thaliana* (2n = 10) (Brassicaceae) has been a model plant across a wide range of research because of its small genome size, short generation time, and the large number of seeds produced from each generation. Because its worldwide distribution covers habitats with stunning ecological complexity throughout Eurasia, Africa, and North America, *A. thaliana* has been used as a model species for revealing the genetic basis and molecular mechanisms for ecological adaptation^1^. The *A. thaliana* genome based on the Col-0 ecotype was completely sequenced and assembled in 2000 as the first plant genome sequence and has greatly promoted molecular studies^2^. With the development of sequencing technology, four versions of *A. thaliana* reference genomes for the same Col-0 ecotype have been reported and updated^3, 4, 5, 6^. Population genomic analyses based on the reference genome and whole-genome re-sequencing data of other ecotypes revealed a widespread global postglacial expansion from sparsely distributed relict ecotypes^7^. In particular, a large number of genetic variations were associated with multiple phenotypic changes underlying ecological adaptation to varied habitats^7^. These genetic variations comprise mainly single nucleotide polymorphisms (SNPs) and short insertions and deletions (INDELs, often < 50 bp) across different ecotypes^8, 9^, as well as allelic variations from major genes associated with ecological phenotypes that have been uncovered through genome-wide association studies (GWAS)^10, 11^.

Beyond SNPs and INDELs, it remains unknown whether both variable genes and large structural variations (SVs, often > 50 bp) contribute to ecological adaptation. The SVs mainly comprise presence/absence variants (PAVs), inversions (INVs), translocations (TRANs), and copy number variations (CNVs), which not only affect gene expression but also remove and produce new genes. SVs were expected and partially confirmed to be important contributors to phenotypic variations^12, 13^. In particular, incomplete detection of genomic variants may lead to weak linkage disequilibrium (LD), which decreases the statistical power of GWAS analyses that ultimately fail to identify the major genetic loci underlying ecological phenotypes^14, 15^. Assembling the high-quality *de novo* genomes of multiple ecotypes^16^ and conducting pan-genome analyses^17, 18^ of these genomes could reveal SVs and capture missing heritability^15^. In addition, a graph-based pan-genome assembly can efficiently integrate genetic variants of all *de novo* genomes and identify the major SVs underlying diverse phenotypes^19, 20, 21^.

In this study, we assembled 38 high-quality genomes for representative ecotypes of *A. thaliana* using PacBio-HiFi long-read sequencing across its distribution. We sampled six sparsely districted relict ecotypes, one from the Qinghai-Tibet Plateau (QTP) of western China, one from Italy, and four from Morocco^22^. The other 32 ecotypes derive from postglacial expansions based on whole-genome re-sequencing analyses of the *Arabidopsis thaliana* 1001 Genome Project (https://1001genomes.org/)^7^ **(Supplementary Table 1)**. These ecotypes cover most major clades and subclades of the 1,135 global accessions and represent the diverse habitats occupied by *A. thaliana*^7^. Based on the assembled high-quality *de novo* genomes, we aimed to address the following questions: (1) how many more genes could be annotated compared to the reference genome based on ecotype Col-0? What are the functions of these variable genes? (2) Could the phylogenomic relationships support postglacial expansions of most ecotypes from the sparse relict ecotypes? (3) What are the functions of the SV-overlapped genes? Have these SVs contributed greatly to the local adaptation of this model species across diverse habitats? Through addressing these questions, our study provides a set of high-quality genetic resources to understand the genomic diversity and evolution underlying the ecological adaptation of *A. thaliana* and to functionally test the role of SVs and the presence and absence of genes that may produce special ecological phenotypes.

## Results

### Chromosome-level genome assemblies of 38 ecotypes

In order to obtain the genome diversity across different *Arabidopsis thaliana* ecotypes, we selected 38 representative accessions from Europe, Asia, Africa, and North America, which include 6 relict ecotypes (1 from China-Tibet, 1 from Italy, and 4 from Morocco) for *de novo* genome assemblies **(Fig. 1a and Supplementary Table 1)**. We generated 2.08 - 8.28□Gb (approximately 15-60 X) high-fidelity (HiFi) reads for the 38 accessions **(Supplementary Table 2)** which we then assembled into contigs using hifiasm and anchored onto the five chromosomes using RagTag with the recently published Col-PEK T2T genome as a reference^6^. The final assembly sizes ranged from 129.4 to 144.9 Mb with contig N50 sizes of 5.91□-20.3 Mb **(Supplementary Table 6)**. The completeness of assemblies was evaluated by Benchmarking Universal Single-Copy Orthologs (BUSCO), with completeness scores of 99.0 to 99.3% (single-copy and duplicated) in the chromosome-scale assemblies **(Supplementary Figure 2 and Supplementary Table 6)**. The evaluations indicated high continuity and high completeness of the 38 *A. thaliana* genome assemblies.

**Fig. 1.**
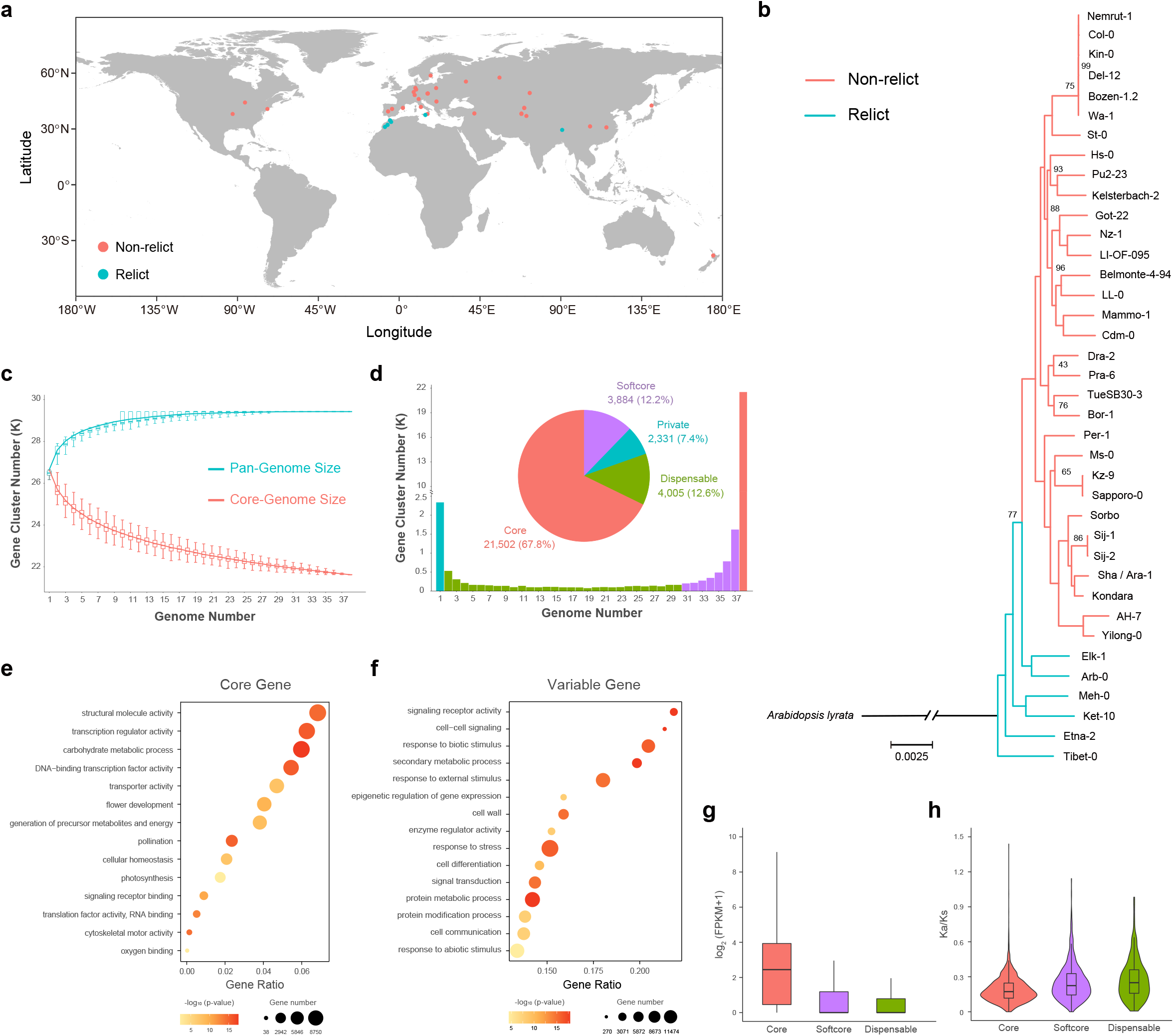
Pan-genome of 38 *A. thaliana* ecotypes. a. Geographic distribution of 38 selected ecotypes of *A. thaliana*. The red circles represent non-relict ecotypes while the green circles represent relict ecotypes. b. Phylogenetic tree of 38 *A. thaliana* ecotypes with *A. lyrata* as outgroup. Branches with 100 bootstrap values are not shown. The red branches represent non-relict ecotypes while the green branches represent relict ecotypes. c. Pan-genome and core genome size simulated by gene cluster number and pan-genome composition. The sample size was set to 1,000 and the sample repeat was set to 30. d. Number and percentage of core, softcore, dispensable, and private gene clusters. e. Bubble chart of GO enrichment analysis for core genes. f. Bubble chart for the GO enrichment analysis of variable genes. g. Expression levels of genes belonging to core, soft-core, and dispensable gene families. h. Pairwise nonsynonymous/synonymous substitution ratios (Ka/Ks) within core, soft-core, and dispensable genes. The upper and lower edges of the boxes represent the 75% and 25% quartiles, the central line denotes the median, and the whiskers extend to 1.5 × IQR in g and h (where IQR is the inter-quartile range).

Combined with transcriptome-based annotation, *ab initio* prediction, homologous-protein-based prediction, and Liftoff mapping using an Araport11 gene annotation file^23^, we finally predicted between 27,239 and 28,823 protein-coding genes in the 38 assembled genomes **(Supplementary Table 7)**. In order to maintain consistency with the Araport11 gene annotation, we prioritized the use of complete gene structures with Liftoff mapping identity and coverage greater than 90%. We also added new annotated genes without overlap, including new genes or genes with structural variations compared with Araport11, including exon shifting. Between 470 and 5,189 genes were found to have structural differences between ecotypes relative to Araport11, and the differences between relict ecotypes were significantly greater than those between non-relict ecotypes **(Supplementary Figure 3**,**4 and Supplementary Table 8)**. The completeness of the gene annotations was also evaluated using BUSCO with completeness scores ranging from 98.9% to 99.7%, suggesting high gene annotation quality **(Supplementary Figure 5 and Supplementary Table 7)**. Around 92.6% to 94.2% of the genes were functionally annotated through at least one of the databases in *eggnog* **(Supplementary Table 7)**.

To infer the evolutionary relationships of the 38 genomes, we clustered the annotated genes into gene families with the sister species *Arabidopsis lyrata* as an outgroup. We selected 17,266 single-copy gene families among these 39 genomes to construct a phylogeny tree. The non-relict accessions clustered into one monophyletic clade while the relict accessions were paraphyletic with the Tibet ecotype which was basal to all other ecotypes **(Fig. 1b)**.

### Pan-genome analyses

We constructed a gene-family-based pan-genome of the 38 ecotypes by clustering a total of 1,059,089 genes into 31,722 pan-gene clusters (including 2,295 clusters with only one gene) using OrthoFinder with the Markov clustering algorithm. Pan-genome size increased with the number of genomes and nearly approached a plateau when n was close to 29 **(Fig. 1c)**. Based on the frequency of occurrence in each genome of the gene clusters, we classified them into four categories: 21,502 (67.8 %) gene families were defined as core gene clusters that were present across all 38 genomes; 3,884 (12.2 %) gene families appeared in 31 to 37 genomes were defined as softcore gene clusters; 4,005 (12.6 %) gene families only found in 2 to 30 genomes were defined as dispensable gene clusters; and 2,331 (7.4 %) accession-specific gene families were defined as private gene clusters **(Fig. 1d)**.

Gene ontology (GO) term enrichment analysis revealed that the core genes were enriched in basic biological and cellular processes, including photosynthesis, cellular homeostasis, and carbohydrate metabolic processes, which suggests that the core genes are mainly involved in maintaining the basic activities of *A. thaliana* **(Fig. 1e)**. Variable genes (including softcore, dispensable and private genes) were enriched in secondary metabolic processes, cell differentiation, and responses to stresses, including responses to biotic/abiotic stimuli and external stimuli **(Fig. 1f)**. In particular, private genes were significantly enriched in response to multiple types of stressors such as endogenous stimuli and light stimuli (**Supplementary Figure 6**). Further investigation of the associations between the presence/absence of variable genes in 38 ecotype genomes and the 19 BIOCLIM environmental variables revealed precipitation seasonality (BIO15) to be significantly associated with the variable genes (**Supplementary Figure 7 and Supplementary Table 9**). Functional enrichment analysis of 525 variable gene families significantly associated with BIO15 showed that these genes were also enriched in response to different types of stress (**Supplementary Figure 8**). These results suggest that the variable genes are likely associated with adaptation to ecotype-specific local environments.

Gene expression analysis showed that the variable genes displayed lower expression levels than the core genes while the non-synonymous/synonymous substitution ratios (Ka/Ks) analysis showed that variable genes had a higher pairwise Ka/Ks value than the core genes **(Fig. 1g, 1h)**. These results suggest that the function of core genes is relatively conservative while variable genes may evolve rapidly to obtain new functions or adapt to the environment.

### The transposable elements (TEs) landscape of 38 *A. thaliana* genomes

We constructed a pan-TE library for the 38 *A. thaliana* genomes based using Repbase and EDTA *de novo* TE annotation and obtained 780 non-redundant TE families (**Supplementary Table 10**). Then we annotated TEs in each genome using RepeatMasker and this pan-TE library and classified the 780 TE families into three categories based on their frequency of occurrence in each genome: core TEs (present in all 38 genomes), variable TEs (present in 6-37 genomes) and rare TEs (present in 1-5 genomes) **(Fig. 2a)**. In all TE families, DNA transposons (26 %) and long terminal repeat-retrotransposons (LTR-RTs) (62 %) accounted for the majority, and variable TEs were mainly LTR type (**Fig. 2b, 2c**). In addition to the TE family differences, the contents of repetitive elements also varied between ecotypes, from 20.34% to 26.44 % of the genomes **(Supplementary Table 6)**. These varied contents led to differences in genome size among ecotypes **(Fig. 2d and Supplementary Figure 9)**. Among all TE categories, LTR-RTs and DNA transposons (such as terminal inverted repeats (TIR)) were the most abundant contents (**Fig. 2e**). We further identified the intact LTR-RTs **(Supplementary Table 11)** and estimated their insertion times into different ecotypes. We found that most of the intact LTR-RTs expanded within the last one million years and numerous LTRs in non-relict ecotypes originated more recently **(Fig. 2f and Supplementary Figure 10)**. This may have led to the emergence of new LTR families and the variety of LTR families between relict and non-relict ecotypes.

**Fig. 2.**
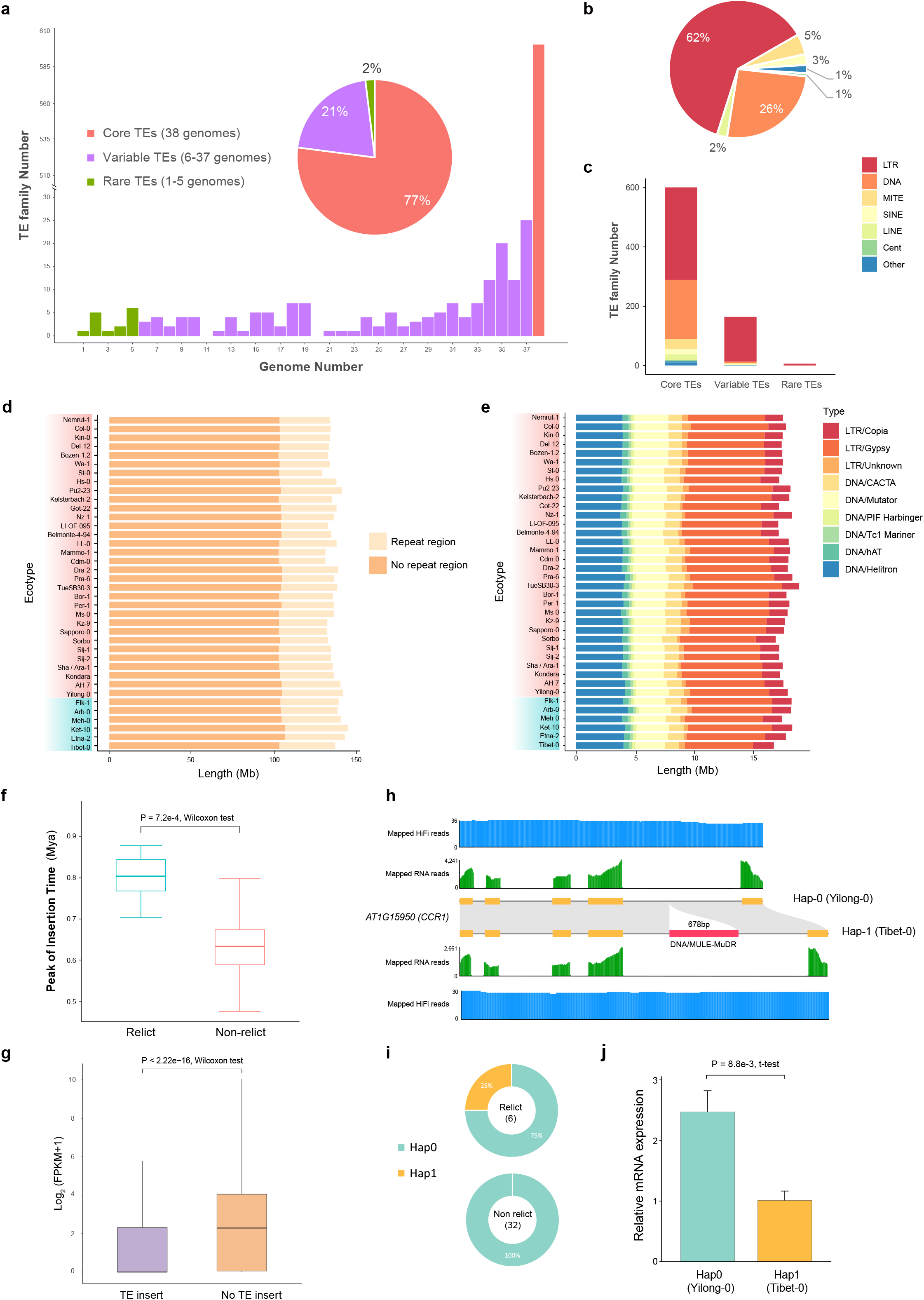
Repetitive sequences of 38 *de novo* genomes. a. Number and percentage of core, variable, and rare TE families. b. Classification of 780 pan-TE families. c. Distribution of TE types in core, variable, and rare TE families. d. TE length identified in different *A thaliana* genomes. e. Composition of different TE types in *A. thaliana* genomes. Blue rectangles display relict ecotypes while red rectangles display non-relict ecotypes. f. Comparison of peak intact LTR-RTs insertion times of relict ecotypes and non-relict ecotypes. Significance was determined using a Wilcoxon test with *p* = 7.2e-4 < 0.05. e. Comparison of the expression levels between genes with and without TE insertion. Significance was determined using a Wilcoxon test with *p* < 2.2e-16 < 0.05. The middle line of the boxplot is the median, the lower and upper hinges correspond to the first and third quartiles, and the upper whisker extends from the hinge to the largest value no further than 1.5 × IQR from the hinge (where IQR is the inter-quartile range), the lower whisker extends from the hinge to the smallest value at most 1.5 × IQR of the hinge, and the outliers are removed in f and g. h. The two haplotypes of *CCR1* are determined by the presence or absence of DNA/MULE-MuDR insertion (red bar) in the fourth intron. HiFi and RNA-seq read mapping supports the gene structure annotation. i. The distributions of the two *CCR1* haplotypes in relict and non-relict ecotypes. j. Relative *CCR1* mRNA levels assessed by quantitative RT-PCR. Significance was determined using a t-test with *p* = 8.8e-3 < 0.05.

To evaluate the effect of TE insertion on gene expression, we compared the gene expression levels of genes with and without TE insertion. Genes with inserted repetitive sequences displayed lower expression levels (**Fig. 2g**). GO enrichment analysis showed that the TE inserted genes were mainly enriched in cell–cell signaling, lipid metabolic processes, and response to stressors, including biotic and external stimuli **(Supplementary Figure 11)**. For example, *CCR1* (AT1G15950) encodes a cinnamoyl CoA reductase involved in lignin biosynthesis and cell proliferation in leaves. The ccr1 mutants exhibit increased ferulic acid (FeA) which has antioxidant activity and reduces the levels of reactive oxygen species (ROS)^24^. Across 38 ecotypes, we found that a specific DNA/MULE-MuDR insertion occurred in the intron region of *CCR1* in two relict ecotypes (Tibet-0 and Meh-0), the expression levels of which were reduced **(Fig. 2h-2j and Supplementary Figure 12)**. Both ecotypes occur in arid habitats^22, 25^ and the inserted TE that reduces gene expression may have promoted the adaptation of both ecotypes to arid habitats through increasing antioxidant activity while reducing ROS.

We also studied the types and locations of TEs inserted around genes and their influence on gene expression. DNA transposons are the dominant type of TE insertion, followed by LTR. TEs were more likely to insert into the upstream regions of the genes (**Supplementary Figure 13**). This may be related to the regulation of gene expression by promoter region. The expression level of genes with TE insertion in the coding sequence (CDS) region decreased the most, and LTR had the greatest impact on gene expression (**Supplementary Figure 14, 15**). The type and location preference of TE insertion may be related to the differential expression of genes in different ecotypes, which further promotes adaptation to different environments.

### Graph-based pan-genome and structural variations (SVs) identification

To identify structural variations across 38 ecotypes, we constructed a graph pan-genome by integrating variants from the Minimap2 alignment with Col-0 as the reference **(Supplementary Figure 16)**. The graph genome comprised a total of 215.05 Mb sequences with 457,367 nodes and 646,863 edges. Among them, 208,764 non-reference nodes were identified, accounting for 80.68 Mb of the map genome. The specific sequences in each ecotype varied from 56.58 Kb to 8.45 Mb with 174 to 49,675 specific edges to connect them to the reference nodes **(Supplementary Table 12)**. On average, each node spans 0.47 Kb and is connected by 1.41 edges. Based on the sequence of the graph genome, we calculated the pan-genome size and core-genome size with the increased number of genomes added to the graph **(Fig. 3a)**. The pan-genome size increased with the number of genomes added, and plateaued when the number of genomes reached 29.

**Fig. 3.**
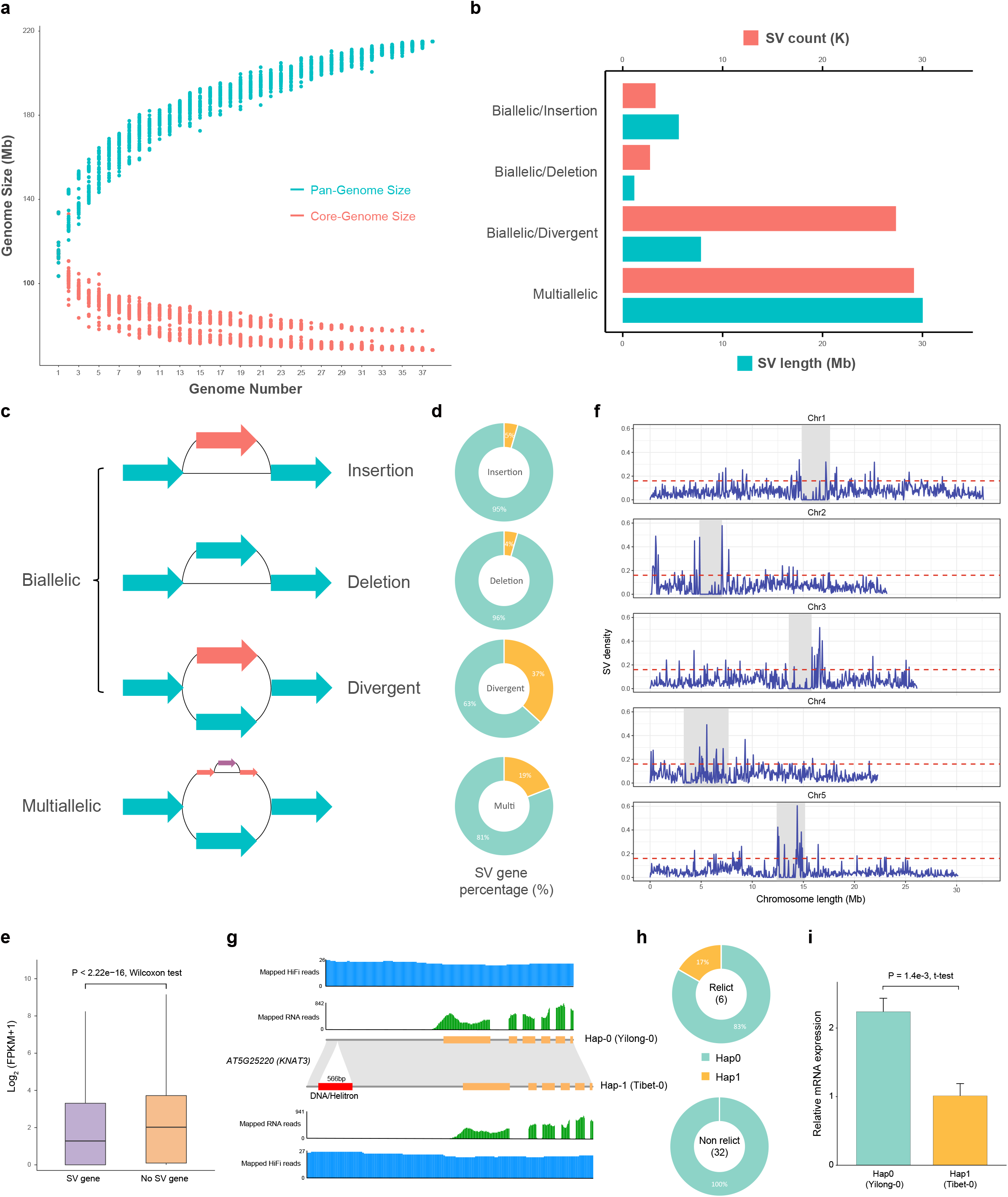
Characterization of the graph genome across 38 *de novo* genomes of *A. thaliana*. a. The graph pan-genome size changes with the increase in number of genome assemblies. b. The bar chart shows the number (red) and length (blue) of each type of SV separately. c. Schematic illustration of diverse SV types from the graph pan-genome based on the reference genome Col-0. d. The pie chart shows the number of genes affected by SV as a proportion of the overall number of genes. e. Expression levels of SV-overlapped genes and no SV-overlapped genes. Significance was determined using a Wilcoxon test with *p* < 2.2e-16 < 0.05. f. SV density along each chromosome based on Col-0 genome assembly: (50 kb sliding windows with a step-size of 20 kb in blue). Gray rectangles: centromeres. The dashed red lines indicate thresholds for SV density values of the top 5%. g. Two haplotypes of *KNAT3* are determined by the presence or absence of DNA/Helitron insertion (red bar) in the promoter region. HiFi and RNA-seq read mapping supports the gene structure annotation. h. The distributions of the two haplotypes of *KNAT3* in relict and non-relict ecotypes. i. Relative *KNAT3* mRNA levels as assessed by quantitative RT-PCR. Significance was determined using a t-test with *p* = 1.4e-3 < 0.05.

We detected structural variations (SVs) in the graph-based genome using gfatools using the bubble popping algorithm and obtained a total of 62,525 SVs shared with at least one genome compared with the reference genome after filtering out SVs with less than 50 bp **(Fig. 3b and Supplementary Table 13)**, and the majority of called SVs were smaller than 500 bp (46,011 out of 62,525, 73.59%) (**Supplementary Figure 17**). SVs were further classified into two types: biallelic (with only one non-reference path) and multiallelic (with more than one non-reference path). The biallelic SVs were further divided into insertion, deletion, and divergent types according to the paths **(Fig. 3c)**. Among the biallelic SVs, the divergent type was the most abundant with a total of 7.82 Mb in length, while the insertion and deletion types only spanned 5.61 Mb and 1.16 Mb, respectively. In addition, the multiallelic type was the largest of the SVs (30.03 Mb), which suggests complex SVs between different ecotypes **(Fig. 3b and Supplementary Table 13)**. Among these SVs, more than 13,924 (22.27 %) were correlated with the inserted TEs and the biallelic/insertion type accounted for the largest proportion (56.9 %) **(Supplementary Table 13)**. Most SVs with inserted TEs were larger than 500bp (8,344 out of 13,924, 59.93%), accounting for 50.53% of total SVs above 500bp, and only 12.12% of total SVs below 500bp had TE insertion. This result suggests that large SVs likely resulted from TE transposition. In addition, the number of SVs was larger in relict ecotypes and there were more SVs specific to relict ecotypes than to non-relict ecotypes derived from postglacial expansion, suggesting distinct differentiations **(Supplementary Figure 18 and Supplementary Table 14)**. The Tibet-0 ecotype had the largest number of specific SVs. However, SVs in non-relict ecotypes had larger TE insertion proportions, which may be related to the more recent TE expansion mentioned above (**Supplementary Table 14**).

We next found the intersection of genes annotated in the Col-0 reference genome with the SV regions. We found 4% to 37% of genes are affected by four types of SVs and the expression levels of these genes were significantly decreased compared to those without SVs **(Fig. 3d, 3e and Supplementary Table 13)**. SVs were more likely to occur in the gene flanking region, and biallelic/divergent SVs affected the largest number of genes (**Supplementary Table 15**). In addition, the expression level of genes with SVs overlapping the CDS region decreased significantly, but the overlapping SVs type had little effect on gene expression **(Supplementary Figure 19, 20)**. Functional enrichment analysis of SVs-overlapped genes showed that they were mainly enriched in secondary metabolic processes, enzyme regulator activity, and responses to diverse stressors, including responses to biotic stimuli and external/endogenous stimuli **(Supplementary Figure 21)**. In addition, GO enrichment results for genes in SV hotspot regions (SV density top 5%) also showed them to be related to catalytic activity and response to light stimuli **(Fig. 3f and Supplementary Figure 22)**. Therefore, the widely distributed variable SVs **(Fig. 3f)** and their overlapped genes may partly account for the ecological adaptation of different ecotypes across diverse habitats.

For example, *KNAT3* (*AT5G25220*) is a class II knotted1-like gene that directly regulates *ABI3* expression with BLH1 to modulate seed germination and early seedling development. The *knat3* mutants are less sensitive to ABA or salinity exposure during seed germination with early seedling development^26^. In addition, *KNAT3* was identified to promote secondary cell wall biosynthesis in xylem vessels together with *KNAT7*. The *knat3 knat7* double mutants had reduced stem tensile and flexural strength compared with wild-type and single mutants^27^. Across 38 ecotypes, we revealed a specific SV in the promoter region of *KNAT3* to the relict Tibet-0 ecotype sampled in the high-altitude Qinghai-Tibet Plateau **(Fig. 3g, h and Supplementary Figure 23)**. Because of this insertion, the gene expression level in Tibet-0 was significantly reduced compared with other ecotypes without this insertion (**Fig. 3i**). The expression level of *KNAT3* was regulated by light as its promoter responded differently to red and far-red light^28^. Therefore, the inserted SV in the *KNAT3* promoter with reduced expression level in Tibet-0 may play an important role in its adaptation to the strong light radiation of the high-altitude region.

### Structural variants (SVs) supplement a large proportion of the heritability and were associated with the variation of multiple environment variables

To test the power of the graph-based pan-genome in capturing missing heritability, we detected 20,326 SVs in 1,073 ecotypes by mapping Illumina short reads to our graph pan-genome. After quality control of minor allele frequency greater than 5%, the remaining SVs were non-randomly distributed across the five chromosomes (**Fig. 4a**), of which only 4,270 (21.01%) were in linkage disequilibrium (LD, **Fig. 4b**) with SNPs. To evaluate the role of SVs in environmental adaptation, we estimated their contribution to the variation of 21 environmental variables (19 BIOCLIM, global UV-B radiation data (https://www.ufz.de/gluv)^29^ and SRTM elevation data from WorldClim v2.1 (www.worldclim.org)^30^) in their natural habitat by fitting a linear mixed model with kinship estimated using SVs. SVs were found to contribute more shares of heritability for 7 (33.33%) environmental variables and explained 78.05% of the phenotypic variations on an average level (**Fig. 4c and Supplementary Table 16**). This is 0.03% more than was explained by SNPs and only 1.36% less than was jointly explained by SV and SNP, indicating an important contribution from SVs as a group to environmental adaptation.

**Fig. 4.**
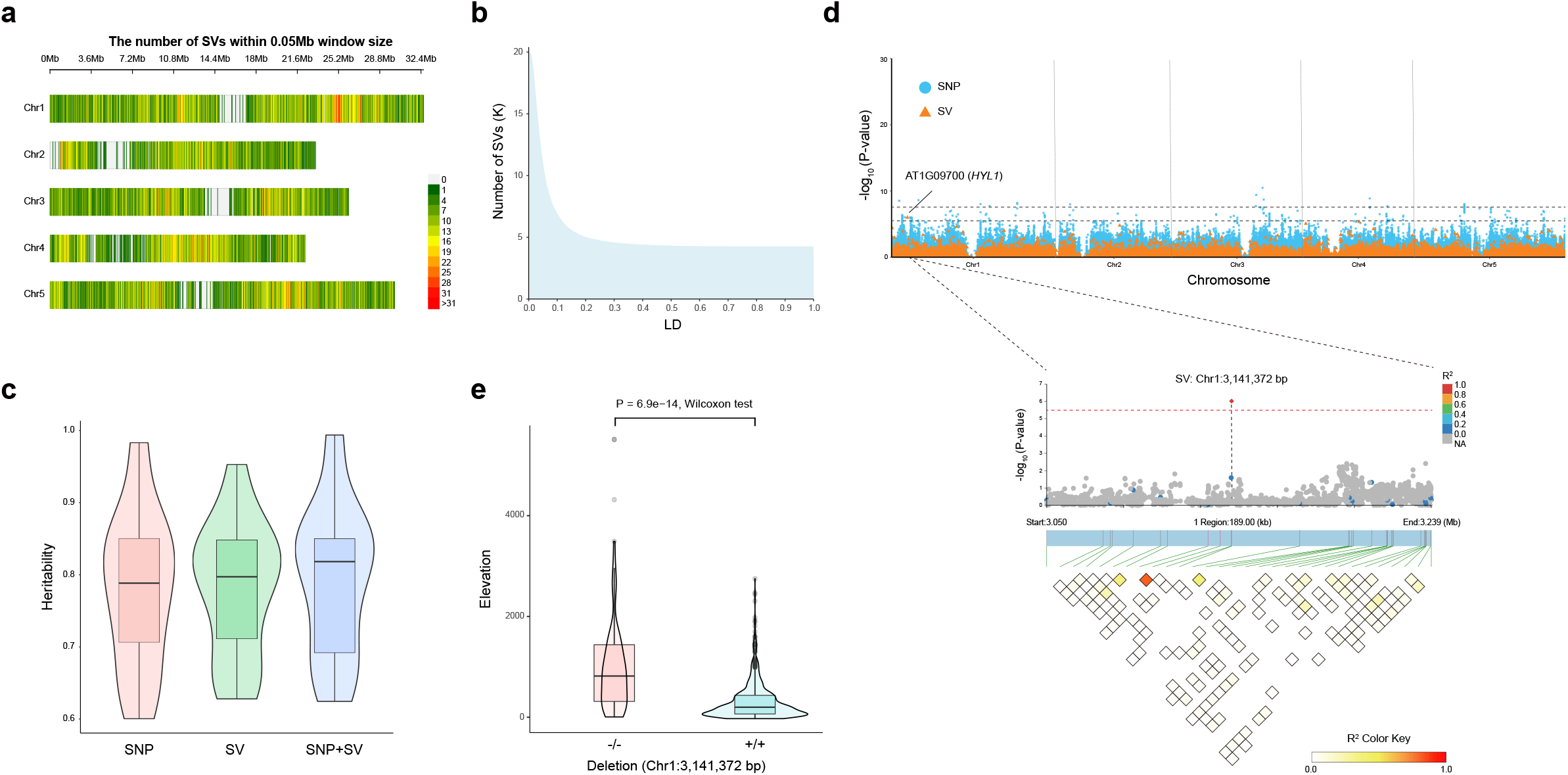
Contribution of structural variants (SVs) to environmental adaptation. a. Genomic distribution of SV from a population of 1,071 worldwide *A. thaliana* accessions from the 1001 Genomes Project (https://1001genomes.org/) and two additional ecotypes, Tibet-0 and Yilong-0. b. Number of SVs (y-axis) tagged by SNPs at a different LD cut-off (x-axis). c. Distribution of heritability (*h*^*2*^) for SNP, SV, and all variants (SV+SNP) estimated from a composite model with SNP and SV. d. Top: Manhattan plot of SV-GWAS (orange) and SNP-GWAS (blue) for elevation. The dashed black lines are genome-wide significance thresholds for SNP-GWAS (upper, 7.55) and SV-GWAS (lower, 5.47). Middle: magnification of the gene region with significant variants is shown and the dot color represents the magnitude of LD (R^2^) with the leading variant SV Chr1:3,141,372. The circles represent SNPs and the triangles represent SVs. Bottom: LD heatmap of the magnified region. e. Boxplot illustrating the genotype and phenotype map at SV associated with elevation. Significance was determined with a Wilcoxon test with *p* = 6.9e-14 < 0.05.

Out of the 21 environmental variables, seven temperature-related variables (BIO1, BIO4, BIO5, BIO6, BIO7, BIO9, BIO11) and elevation showed significant associations in SV-GWAS analyses (**Supplementary Figure 24 and Supplementary Table 17**). For example, one SV peak was associated with elevation variations (**Fig. 4d**). This SV was an 84 bp deletion located in the C-terminal repeat structure of *HYL1* (AT1G09700) of the chromosome 1:3,141,372 bp. *HYL1* was a double-stranded RNA binding protein which is crucial for accurate primary-miRNA processing and the accumulation of mature miRNA (one of the prime regulators of gene expression) in *A. thaliana*, and the C-terminal of *HYL1* regulated its stability and subcellular localization^31^. The mutant *hyl1* was characterized by shorter stature, delayed flowering, leaf hyponasty, reduced fertility, decreased root growth rate, and an altered root gravitropic response^32^. The deletion of this SV might be associated with high-altitude adaptability as it is found in highland ecotypes (-/-) and absent in the lowland ecotypes (+/+) (**Fig. 4e**). Due to a low LD with surrounding SNPs (**Fig. 4d**), no association signals were present in SNP-GWAS around this SV. Taking these results together, the high proportion of variance explained by SVs as a group and the detection of novel SV associations highlighted the value of SV in environment adaptation.

## Discussion

In this study, we assembled high-quality genome sequences of 38 ecotypes in *A. thaliana*. Our phylogenomic analyses of these ecotypes supported the previous hypothesis that *A. thaliana* experienced a postglacial expansion that produced many humid ecotypes across Eurasia and North America^7,23^. These ecotypes comprise a monophyletic lineage despite their widespread distributions. However, six relict ecotypes were paraphyletic and occurred in Europe, Africa, and Asia with long disjunct distances. Interestingly, the Tibet-0 ecotype was inferred to be the earliest-diverged and a sister to the other ecotypes **(Fig. 1b)**. This phylogenomic and phylogeographic pattern suggests that *A. thaliana* should have extended its widespread expansion at least twice from Europe, with the first extending to the QTP where a relict ecotype was retained to the present. Because of the strong selection, this ecotype may have accumulated many specific mutations that clustered it as the earliest divergent ecotype.

In addition to the 67.6 % of the pan-gene-families identified as the core families (21,502 gene families) shared by all ecotypes, the remaining 10,220 gene families (the softcore, dispensable, and private types) vary greatly between ecotypes **(Fig. 1d)**. These variable gene families are functionally enriched in stress responses and associated with climate variables. These findings suggest that gene repertoire varies greatly between ecotypes and gene birth and loss in each ecotype provides a likely basis for local adaptation. This was also confirmed by selection pressures from the core and the variable genes. The core genes have lower Ka/Ks ratios than the variable genes between ecotypes **(Fig. 1h)** and tend to evolve under strong purifying selection^13, 33^. In addition, a total of 62,525 SVs that overlap with 14,243 genes were identified to vary between ecotypes **(Fig. 3b and Supplementary Table 13)**, and these SVs may affect expression levels of the overlapped genes **(Fig. 3e, 3i)**. It should be noted that more than 50% of the identified large SVs (> 500 bp) arise from the inserted TEs. Therefore, it is highly likely that jumping TEs initially created the variable SVs that further removed essential parts of the genes through the reduction of functions, which finally resulted in the polymorphic repertoire with a variable gene number between ecotypes. These genetic changes likely played an important role in the underlying local adaptation of *A. thaliana* to varied habitats.

In addition, using SVs called from 1,135 re-sequencing ecotypes from the *A. thaliana* 1001 Genome Project (https://1001genomes.org/)^7^ and two additional ecotypes, Tibet-0 and Yilong-0, we compared the amount of phenotypic variance explained by SVs and SNPs and found that SVs are an important source of phenotypic variation in addition to SNPs^34^. SVs supplement a large proportion of heritability and are associated with the variation of multiple environmental variables, highlighting their potential contribution to missing heritability and local adaptation^15^. Our genomes, gene annotations, and SVs thus provide valuable resources for systematically exploring the genetic basis underlying how SVs and the deletion and insertion of entire genes contribute to variation in ecological phenotypes and ecological adaptation.

## Methods

### Sample selection and sequencing

We selected 38 representative ecotypes of *Arabidopsis thaliana* distributed on different continents and, using Col-0 as the reference genome, 6 relict *Arabidopsis thaliana* ecotypes, 24 of which had publicly available resequencing data from the *Arabidopsis thaliana* 1001 Genome Project (https://1001genomes.org/)^7^ **(Supplementary Table 1)**. Seeds of the 38 ecotypes were sowed in a greenhouse at Sichuan University until the seeds germinated. Then fresh leaves were collected and stored at −80 °C to construct HiFi SMRTbell libraries. The 15kb libraries were prepared using the SMRTbell Express Template Prep Kit 2.0 (Pacific Biosciences, CA, USA) following the manufacturer’s instructions and sequenced on the PacBio Sequel II platform (Pacific Biosciences, Menlo Park, CA, USA). We used the PacBio SMRT-Analysis package (https://www.pacb.com) for quality control of the raw polymerase reads and generated the HiFi reads by SMRTLink 9.0 software with parameters --min-passes=3 --min-rq=0.99. The final yield HiFi data of 38 accessions ranged from 2.18 Gb to 8.28 Gb, with coverage around 15 to 60 X of the *A. thaliana* genome **(Supplementary Table 2)** based on the k-mer estimate of Col-0 genome size 137.70 Mb as reference **(Supplementary Figure 1 and Supplementary Table 5)**. The total RNA of 12 *A. thaliana* accessions were extracted from the leaf tissues for the library construction. These libraries were subsequently sequenced on the Illumina HiSeq X Ten platform, which produced around 6□Gb of data for each sample **(Supplementary Table 3)**. For whole genome re-sequencing of Tibet-0 and Yilong-0, paired-end libraries were also constructed and sequenced on the Illumina HiSeq X Ten platform **(Supplementary Table 3)**. RNA-seq data of the other 26 accessions were downloaded from the NCBI SRA database under BioProject PRJNA187928^35^, PRJEB15161, and PRJNA319904^36^ **(Supplementary Table 4)**.

### De novo genome assembly of 38 ecotypes

The genome size, heterozygosity, and repeat ratio of the reference Col-0 genome were estimated based on a 17-bp k-mer frequency analysis by GenomeScope v2.0^37^ with parameter ‘-k 17’ and Jellyfish v2.2.9^38^ with parameter ‘-m 17 --min-quality=20 --quality-start=33’ using NGS data download from CRA004538^5^ in CNCB database. Genomes of the 38 sequenced accessions were assembled by hifiasm v 0.18 (https://github.com/chhylp123/hifiasm)^39^ using CCS reads, with parameters ‘-l0’ to disable duplication purging, which may introduce misassemblies if a species has low heterozygosity. There are two outputs of raw hifiasm assemblies: the primary assembly (p_ctg) and the alternate assembly (a_ctg), we selected p_ctg for further assembly and downstream analyses. In order to construct 5 pseudo-chromosomes of each *A. thaliana* accession, we used RagTag v 2.1.0^40^ to scaffold the contigs based on recently published telomere-to-telomere (T2T) genome assembly Col-PEK^6^. The completeness of each assembly was estimated using the embryophyta_odb10 database by Benchmarking Universal Single-Copy Orthologs (BUSCO) v.5.0.2^41^ with default parameters.

### Identification and annotation of repetitive elements

To structurally annotate transposable elements (TEs) in 38 assembled genomes, we used the Extensive De-Novo TE Annotator (EDTA) v.2.1.0^42^ with parameter ‘--species others --sensitive 1 --step all --anno 1 --u 7e-9’ to generate the non-redundant de novo TE libraries and annotated the intact long terminal repeat retrotransposons (LTR-RTs) for each accession. The insertion time of each intact LTR was also provided. The generated TE libraries and *Arabidopsis* repeats in RepBase were further passed into pan-EDTA^43^ to generate the pan-TE library. Repeat regions of 38 genomes were re-masked by RepeatMasker v 4.1.2-p1 (http://www.repeatmasker.org)^44^ with default parameter using pan-TE library. For overlapping repeats, the overlapped regions were split in the middle. Furthermore, to estimate the repetitive elements continuity of each assembly, the LTR assembly index (LAI) was calculated by LTR_retriever v 2.8^45^ using intact LTR datasets.

### Prediction of protein-coding genes

In order to obtain high-quality gene structure annotation of each accession, we combined three different methods: ab initio, protein homology, and transcriptome-based annotation. Firstly, we aligned RNA-seq reads to each genome using HISAT2 v 2.1.1^46^ with parameter ‘--dta’ and assembled transcripts using StringTie v 2.1.4^47^ with parameter ‘--rf’. The assembled transcripts were then passed to PASA v.2.3.3^48^ after filtering by seqclean to generate Open Reading Frames (ORFs). The predicted complete, multi-exon genes models were removed redundant high identity (with an all-to-all identity cut off of 70%) subsequently and sent to train the Hidden Markov Model (HMM) for AUGUSTUS v 3.2.3^49^. In order to further support gene annotation by AUGUSTUS, we also used bam2hints from AUGUSTUS to generate an intron hints file based on a bam file generated by HISAT2. We used this hints file to carry out ab initio gene prediction by AUGUSTUS using default parameters. For homologous prediction protein sequences of Araport11^23^ were downloaded from https://www.arabidopsis.org/ and aligned against each genome using TBLASTN^50^ with parameters ‘-e 1e-5’. After filtering low-quality results, the gene structure was predicted using GeneWise v 2.4.1^51^. The results of PASA, AUGUSTUS, and GeneWise were combined using EvidenceModeler (EVM) v 1.1.1^48^ to generate a combined protein-coding gene set.

As for the model plant, gene numbers starting with ATXG are widely used in *A. thaliana*. In order to minimize the difference from previous gene annotations, we use Liftoff v 1.6.3^52^ to map Araport11 gene annotation onto each genome with parameter ‘-exclude_partial -a 0.9 -s 0.9 -polish’ and replaced our gene annotation which overlaps with the Araport11 gene (valid_ORF=True). The final gene set was named such as col_AT1G01010 (mapped by Araport11) and col00072 (unmapped or newly annotated). The longest transcript of each predicated gene model was considered as the representative for further analysis. The completeness of gene annotations was also estimated by BUSCO using the embryophyta_odb10 database with default parameters.

For gene functional annotation, eggNOG-mapper v2^53^ was applied to get seed ortholog and function description, Gene Ontology (GO) number, Enzyme Commission nomenclature (EC) number, Kyoto Encyclopedia of Genes and Genomes (KEGG) number, PFAM number and so on.

### Phylogenetic analysis

In order to construct phylogenetic relationships among 38 ecotypes of *A. thaliana*, protein sequences from *Arabidopsis lyrata* were downloaded from Phytozome v13 (https://phytozome-next.jgi.doe.gov/)^54^ as an outgroup. Then we did all-to-all blastp with peptide sequences of protein-coding genes annotated from these 39 genomes by NCBI BLAST v 2.2.30+^50^ with cut-off e-values of 1e-5 and input the results into OrthoFinder^55^ for gene clustering with parameter ‘-I 1.5’. The single-copy orthologous genes were further extracted from OrthoFinder results, protein sequences were aligned by MAFFT v 7.490^56^ and extract conserved sites from multiple sequence alignment by Gblocks v 0.91b^57^. The phylogenetic tree was constructed by IQ-TREE v 2.0.3^58^ with parameter ‘-m MFP -B 1000 --bnni’ to automatically find the best model and perform 1000 ultrafast bootstrap analyses to test the robustness of each branch.

### Construction of the protein-coding gene-based pan-genome

We did all-to-all blastp with protein sequences of 38 *A. thaliana* accessions with parameter ‘-e 1e-5’ and input the result file into OrthoFinder for gene family construction by setting the inflation factor to 1.5. Finally, we got 31,722 non-redundant gene clusters. We then classified those clusters into 4 categories: core gene clusters that were conserved in all 38 ecotypes; soft-core gene clusters, which were present in 31–37 accessions; dispensable gene clusters, which were found in 2–30 genomes; and private gene clusters, which contained genes from only 1 sample (including unassigned genes). The longest encoded protein was chosen to represent each gene. In order to further simulate pan-genome size and core genome size by the number of protein-coding genes, we used PanGP v 1.0.1^59^ based on OrthoFinder result with a completely random algorithm setting sample size to 1000 and sample repeat to 30.

### Identification of environment-associated variable gene families

Environmental data from 1970 to 2000 for the 19 BIOCLIM variables were downloaded from WorldClim v2.1 (www.worldclim.org)^30^ with 30 seconds (∼1 km^2^) spatial resolution. Principal component analysis (PCA) of 38 *A. thaliana* ecotypes based on variable gene families was performed by function rda() in R package vegan^60^. Multiple regression of 19 BIOCLIM variables on selected ordination axes was performed by function env.fit() in R package vegan with significance determined using 99,999 permutations. Variable gene families which were significantly associated with BIOCLIM variables were further identified by logistic modeling using the glm() function with parameter ‘family = “binomial”‘.

### Gene expression analysis

We first removed the adaptor sequences and discarded the low-quality reads using Trimmomatic v 0.38^61^ with parameter values ‘SE ILLUMINACLIP:TruSeq3-SE.fa:2:30:10 LEADING:3 TRAILING:3 SLIDINGWINDOW:4:15 MINLEN:36 TOPHRED33’ for single-end RNA-seq reads and ‘PE ILLUMINACLIP:TruSeq3-PE.fa:2:30:10 LEADING:3 TRAILING:3 SLIDINGWINDOW:4:15 MINLEN:36 TOPHRED33’ for paired-end RNA-seq reads. Then the clean reads were mapped to the reference genomes using HISAT2. The expression levels of each gene were calculated using StringTie in terms of FPKM (fragments per kilobase of exon model per million mapped fragments) with the default parameters.

### Ka/Ks calculation of different types of pan-genes

Non-synonymous substitution rates (Ka), synonymous substitution rates (Ks), and Ka/Ks in core, softcore and dispensable gene clusters were computed using the KaKs_Calculator v 2.0^62^ with default parameters. We conducted amino acid alignment for gene pairs in each cluster first and then converted the results into the coding sequence (CDS) alignment using PAL2NAL v 14^63^. The alignments were further passed to the KaKs_Calculator to obtain Ka/Ks values.

### Construction of the graph-based pan-genome and SVs calling

We used minigraph^64^ to construct the graph pangenome of the 38 high-quality *A. thaliana* genome assemblies based on sequence alignment using a modified minimap2 with the parameter ‘-cxggs’. The Col-0 genome assembled in this study was set as the reference and the other 37 genomes were added into the multi-assembly graph successively. The fragments differing from the reference genome are displayed as different paths in the generated graphical fragment assembly (GFA) file. If two or more paths are connected between two fragments, they will form bubbles.

The minigraph graph consists of chains of bubbles with the reference sequences as the backbone. Each bubble in the graph represents a structural variation. In order to call structural variations based on bubbles, we used gfatools (https://github.com/lh3/gfatools) to get the position of a bubble/variation. Extracted structural variations were further classified into biallelic (two paths in a bubble) and multiallelic (more than two paths in a bubble) types.

To genotype the structural variations (SVs) in the 1,135 individuals downloaded from the NCBI SRA database under BioProject PRJNA273563 with 2 individuals, Tibet-0 and Yilong-0, sequenced in this study, we mapped the short reads from each individual to the graph-based pan-genome via vg toolkit v1.40.0-88-g04775076b^19^ using default parameters. After filtering individuals with a missing rate above 0.5 or minor allele frequency (MAF) above 0.05 using plink v1.90b6.7^65^, a total of 1,073 individuals with 20,326 SVs were passed. Then these SVs were imputed using beagle.22Jul22.46e^66^.

### Structural variation gene identification and verification

In order to obtain the actual chromosome position and gene region overlap of SV in different ecotypes, we conducted a whole-genome alignment of 38 *A. thaliana* genomes. The 38 HiFi assembled genomes were aligned to the Col-0 reference genome using Minimap2 v.2.16^67^ with default parameters; alignments lengths shorter than 1,000□bp were discarded. The results show the real positions of each graph pangenome segment in the different genomes and, in combination with the gene annotation files, identified the structural variation genes in the different ecotypes. In order to verify the corresponding relationship between SV genes, we used MCScanX^68^ to perform gene collinearity analysis with default parameters.

In order to confirm the structural variant genes, we mapped HiFi reads to the genome using minimap2 to eliminate assembly errors, while the RNA-seq reads were mapped to the genome using HISAT2 to rule out incorrect gene structure annotations.

### SNP Calling

For SNP database construction, the re-sequencing reads of the 1,135 individuals as well as Tibet-0 and Yilong-0 were mapped to the Col-0 reference genome in this study with the bwa-mem2 algorithm of BWA v0.7.17-r1188^69^ using default parameters, and the resulting BAM files were further filtered using SAMtools v1.3.1^70^ for non-unique and unmapped reads and Picard tools v1.87 (http://broadinstitute.github.io/picard/) for duplication. SNP calling was carried out using the Genome Analysis Toolkit (GATK) v4.2^71^ with default parameters. After filtering via plink with parameters ‘--geno 0.1 --maf 0.05 --mind 0.1’, a total of 2,514,572 SNPs were used for downstream analysis.

### Genome-wide association analysis

We used a standard linear mixed model implemented in GCTA^72^ to perform a genome-wide association analysis for SVs and SNPs. The genetic variants were first filtered by removing alleles with a frequency less than 0.05. A kinship calculated with the genome-wide marker was used to account for confounding with population structure.

### Partitioning the phenotypic variance to SVs and SNPs

We derived a kinship from SV and SNP individually and estimated their heritability by fitting a linear mixed model with the corresponding kinship as a covariance structure implemented in R package hglm^73^. To estimate their joint contribution to phenotypic variance, a composite model with two random effects, one with the SV-derived kinship as the covariance structure and a second with the SNP-derived kinship as the covariance structure, was fitted. Then *h*^*2*^ was estimated as the ratio between the sum of the two variance components from the two random effects and the phenotypic variance.

### RT-qPCR for the *CCR1* and *KNAT3* genes

Total RNA was isolated using the TRIzol method from Tibet-0 and Yilong-0 seedlings. Quality and integrity of the extracted RNA were detected using a NanoDrop 2000 spectrophotometer (Thermo Scientific, Waltham, MA, USA) and 2% agarose gel electrophoresis. We then used the Hifair®III 1st Strand cDNA Synthesis Kit (Yeasen Biotech Co., Ltd, Shanghai, China) to reverse-transcribe the quantified RNA into cDNA. Quantitative Real-time PCR of SV genes was then performed with a Bio-Rad CFX384 Real-Time PCR Detection System (Bio-Rad, USA) using Hifair UNICON Universal Blue qPCR SYBR Green Master Mix (Yeasen Biotech Co., Ltd, Shanghai, China) and the primer sets. The primer sequences used for qRT-PCR analysis are shown in Supplementary Table 18. Each experiment was independently performed three times. Data were normalized to EIF4A by 2^−ΔΔCT^ analysis.

## Data availability

The raw sequencing data for the PacBio HiFi reads, RNA sequencing reads, and re-sequencing Illumina short reads have been deposited in the Genome Sequence Archive (GSA)^74^ database at the National Genomics Data Center, Beijing Institute of Genomics, Chinese Academy of Sciences / China National Center for Bioinformation under BioProject PRJCA012695. The genome assembly and genome annotation have been deposited in the Figshare database (https://dx.doi.org/10.6084/m9.figshare.21673895).

## Acknowledgments

The work was supported by the Natural Science Foundation of China (32030006) and the Second Tibetan Plateau Scientific Expedition and Research (STEP) program (2019QZKK0502).

## Author Contributions

J. L. led the research. S. L. and Y. Z. co-directed the program. S. L. prepared materials. M. K., H. W., W. L., M. Z., Y. H., W. L. and C. C. performed the bioinformatics analysis. K. Y., Y. Z., Z. Y. and H. L. performed the experiments. M. K. wrote the manuscript. M. K., S. L., Y. Z. and J. L. were for writing-review and editing. All authors discussed the results and commented on the manuscript.

## References

1. Provart NJ, et al. 50 years of Arabidopsis research: highlights and future directions. New Phytol 209, 921–944 (2016).

2. AGI. Analysis of the genome sequence of the flowering plant Arabidopsis thaliana. Nature 408, 796–815 (2000).

3. Lamesch P, et al. The Arabidopsis Information Resource (TAIR): improved gene annotation and new tools. Nucleic Acids Res 40, D1202–D1210 (2012).

4. Naish M, et al. The genetic and epigenetic landscape of the Arabidopsis centromeres. Science 374, eabi7489 (2021).

5. Wang B, et al. High-quality Arabidopsis thaliana genome assembly with nanopore and HiFi long reads. Genomics, Proteomics & Bioinformatics 20, 4–13 (2022).

6. Hou X, Wang D, Cheng Z, Wang Y, Jiao Y. A near-complete assembly of an Arabidopsis thaliana genome. Molecular Plant 15, 1247–1250 (2022).

7. Alonso-Blanco C, et al. 1,135 genomes reveal the global pattern of polymorphism in Arabidopsis thaliana. Cell 166, 481–491 (2016).

8. Durvasula A, et al. African genomes illuminate the early history and transition to selfing in Arabidopsis thaliana. Proceedings of the National Academy of Sciences 114, 5213–5218 (2017).

9. Fulgione A, Koornneef M, Roux F, Hermisson J, Hancock AM. Madeiran Arabidopsis thaliana reveals ancient long-range colonization and clarifies demography in Eurasia. Mol Biol Evol 35, 564–574 (2018).

10. Aranzana MJ, et al. Genome-wide association mapping in Arabidopsis identifies previously known flowering time and pathogen resistance genes. PLoS Genet 1, e60 (2005).

11. Atwell S, et al. Genome-wide association study of 107 phenotypes in Arabidopsis thaliana inbred lines. Nature 465, 627–631 (2010).

12. Kosugi S, Momozawa Y, Liu X, Terao C, Kubo M, Kamatani Y. Comprehensive evaluation of structural variation detection algorithms for whole genome sequencing. Genome Biol 20, 1–18 (2019).

13. Göktay M, Fulgione A, Hancock AM. A new catalog of structural variants in 1,301 A. thaliana lines from Africa, Eurasia, and North America reveals a signature of balancing selection at defense response genes. Mol Biol Evol 38, 1498–1511 (2021).

14. Eichler EE, et al. Missing heritability and strategies for finding the underlying causes of complex disease. Nat Rev Genet 11, 446–450 (2010).

15. Zhou Y, et al. Graph pangenome captures missing heritability and empowers tomato breeding. Nature, 1–8 (2022).

16. Hufford MB, et al. De novo assembly, annotation, and comparative analysis of 26 diverse maize genomes. Science 373, 655–662 (2021).

17. Golicz AA, Batley J, Edwards D. Towards plant pangenomics. Plant Biotechnol J 14, 1099–1105 (2016).

18. Dutilh B, Consortium CP-G. Computational pan-genomics: status, promises and challenges. Briefings in Bioinformatics 19, 1 (2018).

19. Garrison E, et al. Variation graph toolkit improves read mapping by representing genetic variation in the reference. Nat Biotechnol 36, 875–879 (2018).

20. Rakocevic G, et al. Fast and accurate genomic analyses using genome graphs. Nat Genet 51, 354–362 (2019).

21. Sirén J, et al. Pangenomics enables genotyping of known structural variants in 5202 diverse genomes. Science 374, abg8871 (2021).

22. Toledo B, Marcer A, Méndez-Vigo B, Alonso-Blanco C, Picó FX. An ecological history of the relict genetic lineage of Arabidopsis thaliana. Environ Exp Bot 170, 103800 (2020).

23. Cheng CY, Krishnakumar V, Chan AP, Thibaud□Nissen F, Schobel S, Town CD. Araport11: a complete reannotation of the Arabidopsis thaliana reference genome. Plant J 89, 789–804 (2017).

24. Xue J, et al. CCR1, an enzyme required for lignin biosynthesis in Arabidopsis, mediates cell proliferation exit for leaf development. Plant J 83, 375–387 (2015).

25. Lou S, et al. Allelic shift in cis-elements of the transcription factor RAP2. 12 underlies adaptation associated with humidity in Arabidopsis thaliana. Sci Adv 8, eabn8281 (2022).

26. Kim D, Cho Yh, Ryu H, Kim Y, Kim TH, Hwang I. BLH 1 and KNAT 3 modulate ABA responses during germination and early seedling development in Arabidopsis. Plant J 75, 755–766 (2013).

27. Wang S, Yamaguchi M, Grienenberger E, Martone PT, Samuels AL, Mansfield SD. The Class II KNOX genes KNAT3 and KNAT7 work cooperatively to influence deposition of secondary cell walls that provide mechanical support to Arabidopsis stems. The Plant J 101, 293–309 (2020).

28. Serikawa KA, Martinez□Laborda A, Kim HS, Zambryski PC. Localization of expression of KNAT3, a class 2 knotted1□like gene. Plant J 11, 853–861 (1997).

29. Beckmann M, et al. gl UV: a global UV□B radiation data set for macroecological studies. Methods Ecol Evol 5, 372–383 (2014).

30. Fick SE, Hijmans RJ. WorldClim 2: new 1□km spatial resolution climate surfaces for global land areas. Int J Climatol 37, 4302–4315 (2017).

31. Bhagat PK, Verma D, Singh K, Badmi R, Sharma D, Sinha AK. Dynamic Phosphorylation of miRNA Biogenesis Factor HYL1 by MPK3 Involving Nuclear–Cytoplasmic Shuttling and Protein Stability in Arabidopsis. Int J Mol Sci 23, 3787 (2022).

32. Lu C, Fedoroff N. A mutation in the Arabidopsis HYL1 gene encoding a dsRNA binding protein affects responses to abscisic acid, auxin, and cytokinin. The Plant Cell 12, 2351–2365 (2000).

33. Zhang L, Li W-H. Mammalian housekeeping genes evolve more slowly than tissue-specific genes. Mol Biol Evol 21, 236–239 (2004).

34. Audano PA, et al. Characterizing the major structural variant alleles of the human genome. Cell 176, 663-675. e619 (2019).

35. Schmitz RJ, et al. Patterns of population epigenomic diversity. Nature 495, 193–198 (2013).

36. Kawakatsu T, et al. Epigenomic diversity in a global collection of Arabidopsis thaliana accessions. Cell 166, 492–505 (2016).

37. Ranallo-Benavidez TR, Jaron KS, Schatz MC. GenomeScope 2.0 and Smudgeplot for reference-free profiling of polyploid genomes. Nat Commun 11, 1–10 (2020).

38. Marçais G, Kingsford C. A fast, lock-free approach for efficient parallel counting of occurrences of k-mers. Bioinformatics 27, 764–770 (2011).

39. Cheng H, Concepcion GT, Feng X, Zhang H, Li H. Haplotype-resolved de novo assembly using phased assembly graphs with hifiasm. Nat Methods 18, 170–175 (2021).

40. Alonge M, et al. Automated assembly scaffolding elevates a new tomato system for high-throughput genome editing. bioRxiv (2021).

41. Simão FA, Waterhouse RM, Ioannidis P, Kriventseva EV, Zdobnov EM. BUSCO: assessing genome assembly and annotation completeness with single-copy orthologs. Bioinformatics 31, 3210–3212 (2015).

42. Ou S, et al. Benchmarking transposable element annotation methods for creation of a streamlined, comprehensive pipeline. Genome Biol 20, 1–18 (2019).

43. Ou S, et al. Differences in activity and stability drive transposable element variation in tropical and temperate maize. bioRxiv (2022).

44. Chen N. Using Repeat Masker to identify repetitive elements in genomic sequences. Curr Protoc in Bioinformatics 5, 4.10. 11-14.10. 14 (2004).

45. Ou S, Jiang N. LTR_retriever: a highly accurate and sensitive program for identification of long terminal repeat retrotransposons. Plant Physiol 176, 1410–1422 (2018).

46. Kim D, Langmead B, Salzberg SL. HISAT: a fast spliced aligner with low memory requirements. Nat Methods 12, 357–360 (2015).

47. Pertea M, Pertea GM, Antonescu CM, Chang T-C, Mendell JT, Salzberg SL. StringTie enables improved reconstruction of a transcriptome from RNA-seq reads. Nat Biotechnol 33, 290–295 (2015).

48. Haas BJ, et al. Automated eukaryotic gene structure annotation using EVidenceModeler and the Program to Assemble Spliced Alignments. Genome Biol 9, 1–22 (2008).

49. Stanke M, Keller O, Gunduz I, Hayes A, Waack S, Morgenstern B. AUGUSTUS: ab initio prediction of alternative transcripts. Nucleic Acids Res 34, W435–W439 (2006).

50. Camacho C, et al. BLAST+: architecture and applications. BMC Bioinformatics 10, 1–9 (2009).

51. Birney E, Clamp M, Durbin R. GeneWise and genomewise. Genome Res 14, 988–995 (2004).

52. Shumate A, Salzberg SL. Liftoff: accurate mapping of gene annotations. Bioinformatics 37, 1639–1643 (2021).

53. Cantalapiedra CP, Hernández-Plaza A, Letunic I, Bork P, Huerta-Cepas J. eggNOG-mapper v2: functional annotation, orthology assignments, and domain prediction at the metagenomic scale. Mol Biol Evol 38, 5825–5829 (2021).

54. Goodstein DM, et al. Phytozome: a comparative platform for green plant genomics. Nucleic Acids Res 40, D1178–D1186 (2012).

55. Emms DM, Kelly S. OrthoFinder: phylogenetic orthology inference for comparative genomics. Genome Biol 20, 1–14 (2019).

56. Katoh K, Standley DM. MAFFT multiple sequence alignment software version 7: improvements in performance and usability. Mol Biol Evol 30, 772–780 (2013).

57. Castresana J. Selection of conserved blocks from multiple alignments for their use in phylogenetic analysis. Mol Biol Evol 17, 540–552 (2000).

58. Minh BQ, et al. IQ-TREE 2: new models and efficient methods for phylogenetic inference in the genomic era. Mol Biol Evol 37, 1530–1534 (2020).

59. Zhao Y, et al. PanGP: a tool for quickly analyzing bacterial pan-genome profile. Bioinformatics 30, 1297–1299 (2014).

60. Oksanen J, et al. Package ‘vegan’. Community ecology package, version 2, 1–295 (2013).

61. Bolger AM, Lohse M, Usadel B. Trimmomatic: a flexible trimmer for Illumina sequence data. Bioinformatics 30, 2114–2120 (2014).

62. Wang D, Zhang Y, Zhang Z, Zhu J, Yu J. KaKs_Calculator 2.0: a toolkit incorporating gamma-series methods and sliding window strategies. Genomics, Proteomics & Bioinformatics 8, 77–80 (2010).

63. Suyama M, Torrents D, Bork P. PAL2NAL: robust conversion of protein sequence alignments into the corresponding codon alignments. Nucleic Acids Res 34, W609–W612 (2006).

64. Li H, Feng X, Chu C. The design and construction of reference pangenome graphs with minigraph. Genome Biol 21, 1–19 (2020).

65. Purcell S, et al. PLINK: a tool set for whole-genome association and population-based linkage analyses. Am J Hum Genet 81, 559–575 (2007).

66. Browning BL, Zhou Y, Browning SR. A one-penny imputed genome from next-generation reference panels. Am J Hum Genet 103, 338–348 (2018).

67. Li H. Minimap2: pairwise alignment for nucleotide sequences. Bioinformatics 34, 3094–3100 (2018).

68. Wang Y, et al. MCScanX: a toolkit for detection and evolutionary analysis of gene synteny and collinearity. Nucleic Acids Res 40, e49–e49 (2012).

69. Vasimuddin M, Misra S, Li H, Aluru S. Efficient architecture-aware acceleration of BWA-MEM for multicore systems. In: 2019 IEEE International Parallel and Distributed Processing Symposium (IPDPS)). IEEE (2019).

70. Li H, et al. The sequence alignment/map format and SAMtools. Bioinformatics 25, 2078–2079 (2009).

71. McKenna A, et al. The Genome Analysis Toolkit: a MapReduce framework for analyzing next-generation DNA sequencing data. Genome Res 20, 1297–1303 (2010).

72. Yang J, Lee SH, Goddard ME, Visscher PM. GCTA: a tool for genome-wide complex trait analysis. Am J Hum Genet 88, 76–82 (2011).

73. Rönnegård L, Shen X, Alam M. hglm: A package for fitting hierarchical generalized linear models. The R Journal 2, 20–28 (2010).

74. Chen T, et al. The genome sequence archive family: toward explosive data growth and diverse data types. Genomics, Proteomics & Bioinformatics 19, 578–583 (2021).

