## Supplementary Information for "The pan-genome and local adaptation of *Arabidopsis thaliana*"

**Tables**

**Supplementary Table 1** Summary of the 38 *Arabidopsis thaliana* ecotypes in this study.

|  | **Name** | **CS Number** | **Country** | **Latitude** | **Longitude** | **Altitude** |
| --- | --- | --- | --- | --- | --- | --- |
| Non-Relict | Yilong-0 | - | China | 31.56402 | 106.7062 | 506 |
|  | Bor-1 | CS22590 | Czech Republic | 49.4013 | 16.2326 | 697 |
|  | Bozen-1.2 | CS76358 | Italy | 46.513 | 11.331 | 409 |
|  | Cdm-0 | CS76410 | Spain | 39.73 | -5.74 | 157 |
|  | Got-22 | CS22609 | Germany | 51.5338 | 9.9355 | 310 |
|  | Kin-0 | CS22654 | United States of America | 44.46 | -85.37 | 938 |
|  | Kondara | CS22651 | Tajikistan | 38.48 | 68.49 | 641 |
|  | Kz-9 | CS22607 | Kazakhstan | 49.5 | 73.1 | 283 |
|  | LL-0 | CS22650 | Spain | 41.59 | 2.49 | 329 |
|  | Mammo-1 | CS76365 | Italy | 38.36 | 16.23 | 146 |
|  | Ms-0 | CS22655 | Russian Federation | 55.7522 | 37.6322 | 2,306 |
|  | Nemrut-1 | CS76398 | Turkey | 38.6425 | 42.2394 | 1,114 |
|  | Pra-6 | CS76416 | Spain | 41.05 | -3.54 | 440 |
|  | Pu2-23 | CS22593 | Czech Republic | 49.42 | 16.36 | 2,698 |
|  | Sij-1 | CS76379 | Uzbekistan | 41.45 | 70.05 | 666 |
|  | Sorbo | CS22653 | Tajikistan | 38.35 | 68.48 | 350 |
|  | TueSB30-3 | CS76403 | Germany | 48.53 | 9.06 | 165 |
|  | Hs-0 | CS76145 | Germany | 52.24 | 9.44 | 42 |
|  | LI-OF-095 | CS76165 | United States of America | 40.9447 | -72.8615 | 148 |
|  | Per-1 | CS76210 | Russian Federation | 58 | 56.3167 | 148 |
|  | St-0 | CS76231 | Sweden | 59 | 18 | 25 |
|  | Kelsterbach-2 | CS28382 | Germany | 50.0667 | 8.5333 | 93 |
|  | Del-12 | CS75573 | Serbia | 44.9444 | 21.1828 | 209 |
|  | Nz-1 | CS28578 | New Zealand | -37.7871 | 175.283 | 48 |
|  | Sapporo-0 | CS28724 | Japan | 43 | 141 | 910 |
|  | Dra-2 | CS28214 | Czech Republic | 49.4167 | 16.2667 | 536 |
|  | Belmonte-4-94 | CS76095 | Italy | 42.1167 | 12.4833 | 218 |
|  | Sha / Ara-1 | CS76382 | Afghanistan | 37.29 | 71.3 | 4,058 |
|  | Sij-2 | CS76380 | Uzbekistan | 41.45 | 70.05 | 2,698 |
|  | Wa-1 | CS22644 | Poland | 52.3 | 21 | 80 |
|  | Col-0 | CS76778 | United States of America | 38.3 | -92.3 | 182 |
|  | AH-7 | - | China | 31.19968 | 115.8067 | 518 |
| Relict | Tibet-0 | - | China | 29.68976 | 91.1861 | 4,317 |
|  | Etna-2 | CS76487 | Italy | 37.69 | 14.98 | 1,660 |
|  | Meh-0 | - | Morocco | 33.9561 | -4.0515 | 1,320 |
|  | Elk-1 | - | Morocco | 32.5352 | -6.015 | 1,231 |
|  | Ket-10 | - | Morocco | 34.9608 | -4.6661 | 1,607 |
|  | Arb-0 | - | Morocco | 31.4199 | -7.5262 | 1,097 |

**Supplementary Table 2** Summary of the 38 *Arabidopsis thaliana* ecotypes DNA sequencing data.

| **Name** | **Number of HiFi reads** | **Total data (G**b) | **Reads length N50 (bp)** | **Mean reads length** | **Sequence coverage^1^** (X) |
| --- | --- | --- | --- | --- | --- |
| Yilong-0 | 363,459 | 6.65 | 19,035 | 18,295 | 48.29 |
| Bor-1 | 299,373 | 3.95 | 15,642 | 13,177 | 28.69 |
| Bozen-1.2 | 203,895 | 2.32 | 13,907 | 11,399 | 16.85 |
| Cdm-0 | 164,271 | 2.62 | 16,862 | 15,934 | 19.03 |
| Got-22 | 345,355 | 4.43 | 14,685 | 12,825 | 32.17 |
| Kin-0 | 197,069 | 2.87 | 15,642 | 14,553 | 20.84 |
| Kondara | 341,635 | 5.10 | 15,053 | 14,916 | 37.04 |
| Kz-9 | 326,232 | 4.18 | 14,288 | 12,806 | 30.36 |
| LL-0 | 352,220 | 4.90 | 15,573 | 13,912 | 35.58 |
| Mammo-1 | 173,628 | 2.33 | 15,451 | 13,443 | 16.92 |
| Ms-0 | 389,833 | 5.38 | 14,790 | 13,800 | 39.07 |
| Nemrut-1 | 266,260 | 3.64 | 15,306 | 13,677 | 26.43 |
| Pra-6 | 240,797 | 3.44 | 14,379 | 14,271 | 24.98 |
| Pu2-23 | 352,556 | 5.14 | 14,834 | 14,582 | 37.33 |
| Sij-1 | 296,021 | 3.89 | 15,104 | 13,154 | 28.25 |
| Sorbo | 558,573 | 4.61 | 9,787 | 8,261 | 33.48 |
| TueSB30-3 | 499,105 | 8.22 | 16,966 | 16,465 | 59.69 |
| Hs-0 | 280,189 | 3.63 | 15,007 | 12,955 | 26.36 |
| LI-OF-095 | 317,489 | 3.81 | 14,078 | 12,014 | 27.67 |
| Per-1 | 301,006 | 4.13 | 15,340 | 13,714 | 29.99 |
| St-0 | 194,285 | 3.07 | 16,515 | 15,805 | 22.29 |
| Kelsterbach-2 | 424,029 | 5.53 | 14,572 | 13,039 | 40.16 |
| Del-12 | 161,775 | 2.08 | 14,614 | 12,842 | 15.11 |
| Nz-1 | 400,667 | 5.50 | 15,013 | 13,730 | 39.94 |
| Sapporo-0 | 386,765 | 4.78 | 14,331 | 12,351 | 34.71 |
| Dra-2 | 444,675 | 6.88 | 15,653 | 15,479 | 49.96 |
| Belmonte-4-94 | 242,616 | 3.61 | 15,003 | 14,884 | 26.22 |
| Sha / Ara-1 | 147,138 | 2.18 | 14,961 | 14,839 | 15.83 |
| Sij-2 | 260,428 | 3.74 | 14,578 | 14,379 | 27.16 |
| Wa-1 | 268,508 | 3.85 | 14,505 | 14,324 | 27.96 |
| Col-0 | 237,301 | 4.44 | 19,185 | 18,724 | 32.24 |
| AH-7 | 291,133 | 3.82 | 14,842 | 13,122 | 27.74 |
| Tibet-0 | 349,873 | 5.99 | 17,608 | 17,134 | 43.50 |
| Etna-2 | 849,782 | 6.86 | 9,382 | 8,072 | 49.82 |
| Meh-0 | 902,680 | 7.99 | 10,117 | 8,850 | 58.02 |
| Elk-1 | 857,356 | 8.15 | 10,493 | 9,504 | 59.19 |
| Ket-10 | 936,164 | 8.28 | 12,176 | 8,848 | 60.13 |
| Arb-0 | 821,175 | 7.75 | 10,397 | 9,431 | 56.28 |

^1^Depth was calculated under the estimate of a genome size of 137.70 Mb.

**Supplementary Table 3** Summary of the 12 *Arabidopsis thaliana* ecotypes RNA sequencing data and DNA re-sequencing data for Tibet-0 and Yilong-0.

| **Name** | **Number of clean reads** | **Total data (G**b) | **Reads length (bp)** | **Insert Size (bp)** | **Tissue** | **Library** |
| --- | --- | --- | --- | --- | --- | --- |
| Yilong-0 | 37,664,356 | 5.65 | 2x150 | 380 | Leaf | RNA |
| LI-OF-095 | 37,104,946 | 5.57 | 2x150 | 380 | Leaf | RNA |
| Kelsterbach-2 | 35,826,004 | 5.37 | 2x150 | 380 | Leaf | RNA |
| Del-12 | 34,918,412 | 5.24 | 2x150 | 380 | Leaf | RNA |
| Dra-2 | 35,761,084 | 5.36 | 2x150 | 380 | Leaf | RNA |
| Belmonte-4-94 | 34,019,716 | 5.10 | 2x150 | 380 | Leaf | RNA |
| AH-7 | 33,541,202 | 5.03 | 2x150 | 380 | Leaf | RNA |
| Tibet-0 | 38,939,972 | 5.84 | 2x150 | 380 | Leaf | RNA |
| Meh-0 | 36,202,862 | 5.43 | 2x150 | 380 | Leaf | RNA |
| Elk-1 | 37,121,446 | 5.57 | 2x150 | 380 | Leaf | RNA |
| Ket-10 | 39,128,482 | 5.87 | 2x150 | 380 | Leaf | RNA |
| Arb-0 | 40,749,242 | 6.11 | 2x150 | 380 | Leaf | RNA |
| Yilong-0 | 44,662,924 | 6.70 | 2x150 | 380 | Leaf | DNA |
| Tibet-0 | 60,291,072 | 9.04 | 2x150 | 380 | Leaf | DNA |

**Supplementary Table 4** Summary of the 26 *Arabidopsis thaliana* ecotypes RNA sequencing data downloaded from NCBI SRA database.

| **Name** | **Number of clean reads** | **Total data (G**b) | **Reads length (bp)** | **SRA number** | **Tissue** |
| --- | --- | --- | --- | --- | --- |
| Bor-1 | 50,350,855 | 5.04 | 100 | SRX1734500 | Leaf |
| Bozen-1.2 | 61,075,841 | 6.11 | 100 | SRX1735112 | Leaf |
| Cdm-0 | 27,206,548 | 2.72 | 100 | SRX1735085 | Leaf |
| Got-22 | 19,665,985 | 2.01 | 2x51 | ERX1659469 | Leaf |
| Kin-0 | 31,679,825 | 1.58 | 50 | SRX244062 | Leaf |
| Kondara | 32,249,429 | 1.61 | 50 | SRX244065 | Leaf |
| Kz-9 | 28,682,458 | 1.43 | 50 | SRX244069 | Leaf |
| LL-0 | 22,747,781 | 2.27 | 100 | SRX1734570 | Leaf |
| Mammo-1 | 30,691,722 | 3.07 | 100 | SRX1735104 | Leaf |
| Ms-0 | 23,945,701 | 2.44 | 2x51 | ERX1659521 | Leaf |
| Nemrut-1 | 50,928,062 | 6.09 | 100 / 130 | SRX1735128 | Leaf |
| Pra-6 | 46,299,849 | 4.63 | 100 | SRX1735090 | Leaf |
| Pu2-23 | 27,962,065 | 1.40 | 50 | SRX244087 | Leaf |
| Sij-1 | 8,675,693 | 0.87 | 100 | SRX1734411 | Leaf |
| Sorbo | 14,789,077 | 1.51 | 2x51 | ERX1659578 | Leaf |
| TueSB30-3 | 25,926,685 | 2.59 | 100 | SRX1735133 | Leaf |
| Hs-0 | 27,313,438 | 1.37 | 50 | SRX244052 | Leaf |
| Per-1 | 36,800,151 | 1.84 | 50 | SRX244083 | Leaf |
| St-0 | 16,576,939 | 1.69 | 2x51 | ERX1659588 | Leaf |
| Nz-1 | 26,722,923 | 2.67 | 100 | SRX1734610 | Leaf |
| Sapporo-0 | 32,246,970 | 3.22 | 100 | SRX1734621 | Leaf |
| Sha / Ara-1 | 53,209,654 | 5.32 | 100 | SRX1734418 | Leaf |
| Sij-2 | 33,211,699 | 3.32 | 100 | SRX1734412 | Leaf |
| Wa-1 | 21,672,045 | 1.08 | 50 | SRX244120 | Leaf |
| Col-0 | 74,711,902 | 21.77 | 2x146 | SRX1735134 | Leaf |
| Etna-2 | 61,441,623 | 6.14 | 100 | SRX1734938 | Leaf |

**Supplementary Table 5** K-mer analysis of the Col-0 genome by using K-mer=17.

| **Name** | **K-mer** | **K-mer number** | **K-mer Depth (X)** | **Genome Size (Mb)** | **Heterozygous Ratio (%)** | **Repeat (%)** |
| --- | --- | --- | --- | --- | --- | --- |
| Col-0 | 17 | 6,471,685,116 | 47 | 137.70 | 0.215 | 32.58 |

**Supplementary Table 6** Summary of the 38 *Arabidopsis thaliana* ecotypes genome assembly.

| **Name** | **Genome length (Mb)** | **Contig N50 (Mb)** | **Scaffold N50 (Mb)** | **BUSCO (%)** | **Repeat rate (%)** |
| --- | --- | --- | --- | --- | --- |
| Yilong-0 | 141.8 | 7.35 | 27.56 | 99.2 | 25.99 |
| Bor-1 | 135.7 | 10.72 | 27.16 | 99.2 | 24.07 |
| Bozen-1.2 | 132.9 | 7.88 | 26.04 | 99.3 | 22.68 |
| Cdm-0 | 131.2 | 6.94 | 25.49 | 99.3 | 21.52 |
| Got-22 | 137.9 | 9.06 | 27.07 | 99.3 | 24.40 |
| Kin-0 | 134.0 | 10.87 | 26.12 | 99.3 | 22.72 |
| Kondara | 136.4 | 12.32 | 25.76 | 99.3 | 23.96 |
| Kz-9 | 132.4 | 6.96 | 26.36 | 99.3 | 22.14 |
| LL-0 | 138.0 | 10.61 | 27.35 | 99.2 | 25.24 |
| Mammo-1 | 131.2 | 6.72 | 26.90 | 99.2 | 21.68 |
| Ms-0 | 136.3 | 9.61 | 26.53 | 99.3 | 24.46 |
| Nemrut-1 | 133.8 | 12.17 | 26.15 | 99.2 | 22.79 |
| Pra-6 | 136.6 | 8.22 | 26.10 | 99.2 | 23.25 |
| Pu2-23 | 141.1 | 12.41 | 27.06 | 99.3 | 26.25 |
| Sij-1 | 134.6 | 9.02 | 25.39 | 99.2 | 23.61 |
| Sorbo | 132.7 | 6.27 | 23.30 | 99.1 | 21.87 |
| TueSB30-3 | 138.4 | 20.30 | 26.96 | 99.2 | 24.64 |
| Hs-0 | 137.9 | 7.54 | 26.96 | 99.2 | 24.75 |
| LI-OF-095 | 132.7 | 7.97 | 24.62 | 99.2 | 21.42 |
| Per-1 | 136.4 | 9.87 | 26.74 | 99.2 | 23.29 |
| St-0 | 129.4 | 7.82 | 24.67 | 99.2 | 20.34 |
| Kelsterbach-2 | 135.4 | 13.22 | 27.38 | 99.2 | 24.00 |
| Del-12 | 133.5 | 9.20 | 26.12 | 99.3 | 22.59 |
| Nz-1 | 136.5 | 10.38 | 27.11 | 99.2 | 23.47 |
| Sapporo-0 | 132.1 | 11.10 | 26.39 | 99.2 | 22.08 |
| Dra-2 | 138.9 | 14.61 | 27.08 | 99.3 | 24.55 |
| Belmonte-4-94 | 134.9 | 11.95 | 26.62 | 99.1 | 22.51 |
| Sha / Ara-1 | 135.6 | 6.28 | 26.43 | 99.3 | 23.86 |
| Sij-2 | 134.7 | 7.84 | 26.18 | 99.2 | 23.60 |
| Wa-1 | 134.0 | 13.97 | 26.15 | 99.3 | 22.77 |
| Col-0 | 134.4 | 14.27 | 26.11 | 99.2 | 23.00 |
| AH-7 | 140.3 | 7.52 | 25.53 | 99.2 | 25.42 |
| Tibet-0 | 137.2 | 13.73 | 26.23 | 99.0 | 24.77 |
| Etna-2 | 143.0 | 9.93 | 27.29 | 99.1 | 25.43 |
| Meh-0 | 140.5 | 5.91 | 26.79 | 99.2 | 25.52 |
| Elk-1 | 139.2 | 8.66 | 27.12 | 99.2 | 25.38 |
| Ket-10 | 144.9 | 8.08 | 27.89 | 99.2 | 26.44 |
| Arb-0 | 138.6 | 17.56 | 27.08 | 99.1 | 24.86 |

**Supplementary Table 7** Summary of the 38 *Arabidopsis thaliana* ecotypes gene annotation.

| **Name** | **Gene Number** | **BUSCO** | **Functional annotation number** | **Annotation rate** |
| --- | --- | --- | --- | --- |
| Yilong-0 | 27,683 | 99.4% | 25,916 | 93.62% |
| Bor-1 | 27,801 | 99.3% | 26,087 | 93.83% |
| Bozen-1.2 | 28,738 | 99.7% | 26,684 | 92.85% |
| Cdm-0 | 27,666 | 99.1% | 25,963 | 93.84% |
| Got-22 | 27,835 | 99.2% | 26,066 | 93.64% |
| Kin-0 | 28,744 | 99.6% | 26,711 | 92.93% |
| Kondara | 27,489 | 99.3% | 25,859 | 94.07% |
| Kz-9 | 27,545 | 99.2% | 25,895 | 94.01% |
| LL-0 | 27,778 | 99.2% | 26,066 | 93.84% |
| Mammo-1 | 27,717 | 99.0% | 25,977 | 93.72% |
| Ms-0 | 27,793 | 99.2% | 26,075 | 93.82% |
| Nemrut-1 | 28,752 | 99.5% | 26,678 | 92.79% |
| Pra-6 | 27,772 | 99.3% | 26,002 | 93.63% |
| Pu2-23 | 27,962 | 99.2% | 26,183 | 93.64% |
| Sij-1 | 27,521 | 99.2% | 25,822 | 93.83% |
| Sorbo | 27,646 | 99.1% | 25,913 | 93.73% |
| TueSB30-3 | 27,805 | 99.1% | 26,092 | 93.84% |
| Hs-0 | 27,838 | 99.4% | 26,092 | 93.73% |
| LI-OF-095 | 27,865 | 99.2% | 26,086 | 93.62% |
| Per-1 | 27,793 | 98.9% | 26,019 | 93.62% |
| St-0 | 27,775 | 99.1% | 26,018 | 93.67% |
| Kelsterbach-2 | 27,956 | 99.3% | 26,157 | 93.56% |
| Del-12 | 28,711 | 99.7% | 26,687 | 92.95% |
| Nz-1 | 27,866 | 99.5% | 26,099 | 93.66% |
| Sapporo-0 | 27,598 | 99.2% | 25,924 | 93.93% |
| Dra-2 | 27,913 | 99.3% | 26,138 | 93.64% |
| Belmonte-4-94 | 27,716 | 99.2% | 25,953 | 93.64% |
| Sha / Ara-1 | 27,471 | 99.2% | 25,802 | 93.92% |
| Sij-2 | 27,535 | 99.2% | 25,856 | 93.90% |
| Wa-1 | 28,823 | 99.7% | 26,696 | 92.62% |
| Col-0 | 28,735 | 99.7% | 26,625 | 92.66% |
| AH-7 | 27,531 | 99.3% | 25,867 | 93.96% |
| Tibet-0 | 27,239 | 99.1% | 25,650 | 94.17% |
| Etna-2 | 27,812 | 99.1% | 26,096 | 93.83% |
| Meh-0 | 27,661 | 99.0% | 25,935 | 93.76% |
| Elk-1 | 27,686 | 99.5% | 26,018 | 93.98% |
| Ket-10 | 27,607 | 99.0% | 25,951 | 94.00% |
| Arb-0 | 27,711 | 99.2% | 26,009 | 93.86% |

**Supplementary Table 8** Our gene annotation of 38 *Arabidopsis thaliana* ecotypes’ genome assemblies compare with Araport11 gene annotation.

| **Name** | **Mapped gene number^1^** | **Unmapped gene number** | **New genes or genes with structure variant number** | **Final gene number** |
| --- | --- | --- | --- | --- |
| Yilong-0 | 23,439 | 4,215 | 4,244 | 27,683 |
| Bor-1 | 24,136 | 3,518 | 3,665 | 27,801 |
| Bozen-1.2 | 27,165 | 489 | 1,573 | 28,738 |
| Cdm-0 | 23,903 | 3,751 | 3,763 | 27,666 |
| Got-22 | 24,183 | 3,471 | 3,652 | 27,835 |
| Kin-0 | 27,181 | 473 | 1,563 | 28,744 |
| Kondara | 23,651 | 4,003 | 3,838 | 27,489 |
| Kz-9 | 23,716 | 3,938 | 3,829 | 27,545 |
| LL-0 | 24,043 | 3,611 | 3,735 | 27,778 |
| Mammo-1 | 23,901 | 3,753 | 3,816 | 27,717 |
| Ms-0 | 23,851 | 3,803 | 3,942 | 27,793 |
| Nemrut-1 | 27,174 | 480 | 1,578 | 28,752 |
| Pra-6 | 24,144 | 3,510 | 3,628 | 27,772 |
| Pu2-23 | 24,390 | 3,264 | 3,572 | 27,962 |
| Sij-1 | 23,738 | 3,916 | 3,783 | 27,521 |
| Sorbo | 23,739 | 3,915 | 3,907 | 27,646 |
| TueSB30-3 | 24,131 | 3,523 | 3,674 | 27,805 |
| Hs-0 | 24,157 | 3,497 | 3,681 | 27,838 |
| LI-OF-095 | 24,038 | 3,616 | 3,827 | 27,865 |
| Per-1 | 23,829 | 3,825 | 3,964 | 27,793 |
| St-0 | 24,300 | 3,354 | 3,475 | 27,775 |
| Kelsterbach-2 | 24,404 | 3,250 | 3,552 | 27,956 |
| Del-12 | 27,156 | 498 | 1,555 | 28,711 |
| Nz-1 | 24,176 | 3,478 | 3,690 | 27,866 |
| Sapporo-0 | 23,716 | 3,938 | 3,882 | 27,598 |
| Dra-2 | 24,107 | 3,547 | 3,806 | 27,913 |
| Belmonte-4-94 | 24,123 | 3,531 | 3,593 | 27,716 |
| Sha / Ara-1 | 23,673 | 3,981 | 3,798 | 27,471 |
| Sij-2 | 23,728 | 3,926 | 3,807 | 27,535 |
| Wa-1 | 27,184 | 470 | 1,639 | 28,823 |
| Col-0 | 27,173 | 481 | 1,562 | 28,735 |
| AH-7 | 23,297 | 4,357 | 4,234 | 27,531 |
| Tibet-0 | 22,465 | 5,189 | 4,774 | 27,239 |
| Etna-2 | 23,246 | 4,408 | 4,566 | 27,812 |
| Meh-0 | 23,016 | 4,638 | 4,645 | 27,661 |
| Elk-1 | 23,333 | 4,321 | 4,353 | 27,686 |
| Ket-10 | 22,901 | 4,753 | 4,706 | 27,607 |
| Arb-0 | 23,404 | 4,250 | 4,307 | 27,711 |

^1^Complete genes of >90% identity and coverage with Araport11.

**Supplementary Table 9** Results of multiple regression between variable genes and 19 BIOCLIM environmental variables with the Monte Carlo permutation test.

|  | **PC1** | **PC2** | **R^2^** | **Pr(>r)** |
| --- | --- | --- | --- | --- |
| BIO1 | 0.14348 | 0.98965 | 0.0249 | 0.64377 |
| BIO2 | 0.99988 | 0.0157 | 0.0219 | 0.68217 |
| BIO3 | 0.98409 | -0.17764 | 0.0166 | 0.74438 |
| BIO4 | -0.58149 | -0.81355 | 0.004 | 0.93308 |
| BIO5 | 0.2013 | 0.97953 | 0.0154 | 0.76115 |
| BIO6 | -0.05858 | 0.99828 | 0.0125 | 0.80445 |
| BIO7 | 0.60916 | -0.79305 | 0.0021 | 0.96405 |
| BIO8 | 0.8022 | 0.59706 | 0.0254 | 0.63654 |
| BIO9 | -0.67913 | 0.73402 | 0.0049 | 0.91738 |
| BIO10 | 0.03904 | 0.99924 | 0.0231 | 0.66176 |
| BIO11 | 0.2179 | 0.97597 | 0.0165 | 0.74922 |
| BIO12 | -0.95146 | -0.30777 | 0.0843 | 0.20824 |
| BIO13 | -0.80434 | 0.59417 | 0.0558 | 0.36194 |
| BIO14 | -0.5056 | -0.86277 | 0.1013 | 0.14886 |
| **BIO15** | **0.23676** | **0.97157** | **0.209** | **0.01802*** |
| BIO16 | -0.8444 | 0.53571 | 0.0917 | 0.17805 |
| BIO17 | -0.41044 | -0.91189 | 0.1327 | 0.08052 |
| BIO18 | -0.72075 | 0.6932 | 0.0148 | 0.77056 |
| BIO19 | -0.8948 | -0.44646 | 0.1372 | 0.07483 |

**Supplementary Table 10** Summary of pan-TE library constructed by 38 *Arabidopsis thaliana* ecotypes genome assembly.

| **Type** | **Number of sequences in pan-TE library** |
| --- | --- |
| Cent/centromeric_repeat | 6 |
| DNA/CACTA | 14 |
| DNA/DTA | 3 |
| DNA/DTC | 7 |
| DNA/DTH | 3 |
| DNA/DTM | 9 |
| DNA/DTT | 1 |
| DNA/Harbinger | 3 |
| DNA/Helitron | 61 |
| DNA/Mariner | 7 |
| DNA/MuDR | 58 |
| DNA/hAT | 22 |
| DNA/unknown | 16 |
| LINE/L1 | 12 |
| LINE/unknown | 7 |
| LTR/Copia | 294 |
| LTR/Gypsy | 84 |
| LTR/Ty3 | 66 |
| LTR/unknown | 34 |
| MITE/DTA | 23 |
| MITE/DTC | 1 |
| MITE/DTH | 5 |
| MITE/DTM | 9 |
| SINE/tRNA | 5 |
| SINE/unknown | 15 |
| Satellite/Satellite | 10 |
| rDNA/spacer | 1 |
| repeat/unknown | 4 |
| Total | 780 |

**Supplementary Table 11** Intact LTR-RTs identified in the 38 *Arabidopsis thaliana* ecotypes genome assembly.

| **Name** | **Total intact LTR-RTs searched** | **Copia** | **Gypsy** | **Unknown** | **Whole genome LAI** |
| --- | --- | --- | --- | --- | --- |
| Yilong-0 | 1,498 | 678 | 640 | 180 | 24.92 |
| Bor-1 | 1,554 | 606 | 708 | 240 | 24.52 |
| Bozen-1.2 | 1,506 | 648 | 576 | 282 | 9.29 |
| Cdm-0 | 1,554 | 672 | 684 | 198 | 20.47 |
| Got-22 | 1,476 | 648 | 642 | 186 | 25.75 |
| Kin-0 | 1,476 | 642 | 570 | 264 | 8.49 |
| Kondara | 1,560 | 762 | 612 | 186 | 21.29 |
| Kz-9 | 1,842 | 816 | 624 | 402 | 22.75 |
| LL-0 | 1,794 | 702 | 702 | 390 | 13.30 |
| Mammo-1 | 1,578 | 546 | 774 | 258 | 5.89 |
| Ms-0 | 1,734 | 732 | 696 | 306 | 26.72 |
| Nemrut-1 | 1,524 | 648 | 582 | 294 | 8.85 |
| Pra-6 | 1,716 | 702 | 744 | 270 | 26.59 |
| Pu2-23 | 1,680 | 726 | 684 | 270 | 19.20 |
| Sij-1 | 1,582 | 780 | 598 | 204 | 20.71 |
| Sorbo | 1,528 | 744 | 610 | 174 | 26.01 |
| TueSB30-3 | 2,082 | 702 | 906 | 474 | 24.28 |
| Hs-0 | 1,536 | 672 | 636 | 228 | 18.50 |
| LI-OF-095 | 1,344 | 552 | 630 | 162 | 19.26 |
| Per-1 | 1,884 | 840 | 702 | 342 | 9.08 |
| St-0 | 1,590 | 606 | 708 | 276 | 12.02 |
| Kelsterbach-2 | 1,632 | 600 | 804 | 228 | 14.55 |
| Del-12 | 1,446 | 642 | 576 | 228 | 9.02 |
| Nz-1 | 1,632 | 636 | 720 | 276 | 20.31 |
| Sapporo-0 | 1,818 | 804 | 624 | 390 | 23.05 |
| Dra-2 | 1,632 | 666 | 714 | 252 | 25.89 |
| Belmonte-4-94 | 1,650 | 612 | 750 | 288 | 20.96 |
| Sha / Ara-1 | 1,720 | 900 | 664 | 156 | 25.40 |
| Sij-2 | 1,564 | 780 | 592 | 192 | 25.59 |
| Wa-1 | 1,518 | 648 | 582 | 288 | 8.89 |
| Col-0 | 1,662 | 648 | 582 | 432 | 9.11 |
| AH-7 | 1,426 | 660 | 634 | 132 | 21.02 |
| Tibet-0 | 1,656 | 852 | 504 | 300 | 21.45 |
| Etna-2 | 1,614 | 696 | 642 | 276 | 18.40 |
| Meh-0 | 1,536 | 798 | 558 | 180 | 19.27 |
| Elk-1 | 1,878 | 798 | 756 | 324 | 21.70 |
| Ket-10 | 1,884 | 864 | 798 | 222 | 26.91 |
| Arb-0 | 1,638 | 804 | 618 | 216 | 20.79 |

**Supplementary Table 12** Summary of the graph pan-genome constructed by 38 *Arabidopsis thaliana* ecotype genomes.

| **Name** | **Node number** | **Node total length (bp)** | **Edge number** |
| --- | --- | --- | --- |
| Yilong-0 | 16,545 | 5,635,106 | 30,489 |
| Bor-1 | 2,380 | 1,253,016 | 5,011 |
| Bozen-1.2 | 81 | 56,583 | 174 |
| Cdm-0 | 3,754 | 1,664,225 | 7,942 |
| Got-22 | 15,731 | 5,707,389 | 26,908 |
| Kin-0 | 180 | 74,301 | 381 |
| Kondara | 1,323 | 402,591 | 2,774 |
| Kz-9 | 272 | 114,573 | 500 |
| LL-0 | 4,950 | 2,322,450 | 9,994 |
| Mammo-1 | 5,603 | 2,366,958 | 11,072 |
| Ms-0 | 6,005 | 2,758,970 | 11,215 |
| Nemrut-1 | 104 | 112,107 | 212 |
| Pra-6 | 4,306 | 1,813,301 | 8,130 |
| Pu2-23 | 2,489 | 1,080,101 | 5,210 |
| Sij-1 | 4,412 | 1,698,877 | 8,641 |
| Sorbo | 2,211 | 869,140 | 4,600 |
| TueSB30-3 | 3,185 | 1,407,076 | 6,437 |
| Hs-0 | 3,777 | 1,883,679 | 7,580 |
| LI-OF-095 | 29,576 | 8,453,514 | 49,675 |
| Per-1 | 4,956 | 2,218,627 | 9,721 |
| St-0 | 12,242 | 4,831,755 | 21,374 |
| Kelsterbach-2 | 7,386 | 3,259,330 | 13,705 |
| Del-12 | 105 | 129,278 | 225 |
| Nz-1 | 4,574 | 1,865,168 | 8,929 |
| Sapporo-0 | 8,344 | 3,530,230 | 15,932 |
| Dra-2 | 3,303 | 1,461,896 | 6,689 |
| Belmonte-4-94 | 5,454 | 2,223,777 | 10,796 |
| Sha / Ara-1 | 2,445 | 1,237,019 | 4,830 |
| Sij-2 | 461 | 105,792 | 906 |
| Wa-1 | 179 | 161,814 | 260 |
| **Col-0 (reference)** | 248,573 | 134,371,593 | 248,568 |
| AH-7 | 3,222 | 968,820 | 6,685 |
| Tibet-0 | 8,928 | 3,466,208 | 18,727 |
| Etna-2 | 7,355 | 2,911,800 | 15,367 |
| Meh-0 | 7,106 | 2,718,953 | 14,619 |
| Elk-1 | 6,883 | 2,475,371 | 13,861 |
| Ket-10 | 10,755 | 4,227,870 | 21,734 |
| Arb-0 | 8,482 | 3,212,769 | 16,990 |
| Total | 457,367 | 215,052,507 | 646,863 |

**Supplementary Table 13** Summary of the structural variations (SVs) detected in graph-based pan-genome constructed by 38 *Arabidopsis thaliana* ecotype genomes.

| **SV type** | **SV number** | **SV length (bp)** | **SV influenced gene number** | **SV influenced gene percentage (%)** | **SV with TE inserted** | **SV with TE inserted percentage (%)** |
| --- | --- | --- | --- | --- | --- | --- |
| Biallelic/Insertion | 3,285 | 5,614,354 | 1,325 | 4.61 | 1,870 | 56.9 |
| Biallelic/Deletion | 2,733 | 1,161,514 | 1,273 | 4.43 | 775 | 28.4 |
| Biallelic/Divergent | 27,353 | 7,823,469 | 10,591 | 36.83 | 3,084 | 11.27 |
| Multiallelic | 29,154 | 30,032,134 | 5,469 | 19.02 | 8,195 | 28.11 |

**Supplementary Table 14** Summary of SV classification and TE insertion in the 38 *Arabidopsis thaliana* ecotypes genome assembly.

| **Name** | **Ecotype specific SV** | **Total SV** | **Ecotype specific SV with TE insertion** | **Total SV with TE insertion** |
| --- | --- | --- | --- | --- |
| Yilong-0 | 310 | 12,494 | 81 | 1,688 |
| Bor-1 | 403 | 9,887 | 104 | 1,398 |
| Bozen-1.2 | 11 | 184 | 8 | 79 |
| Cdm-0 | 790 | 10,856 | 162 | 1,411 |
| Got-22 | 402 | 10,149 | 103 | 1,420 |
| Kin-0 | 11 | 167 | 7 | 59 |
| Kondara | 151 | 11,197 | 30 | 1,508 |
| Kz-9 | 1 | 11,310 | 1 | 1,553 |
| LL-0 | 706 | 10,509 | 175 | 1,496 |
| Mammo-1 | 675 | 10,551 | 171 | 1,434 |
| Ms-0 | 453 | 10,051 | 109 | 1,393 |
| Nemrut-1 | 16 | 156 | 10 | 51 |
| Pra-6 | 441 | 9,858 | 110 | 1,439 |
| Pu2-23 | 422 | 8,929 | 82 | 1,281 |
| Sij-1 | 4 | 11,289 | 3 | 1,557 |
| Sorbo | 294 | 11,133 | 76 | 1,557 |
| TueSB30-3 | 426 | 10,060 | 89 | 1,423 |
| Hs-0 | 479 | 9,531 | 87 | 1,293 |
| LI-OF-095 | 423 | 10,139 | 107 | 1,456 |
| Per-1 | 568 | 11,003 | 122 | 1,519 |
| St-0 | 454 | 9,292 | 124 | 1,374 |
| Kelsterbach-2 | 457 | 9,158 | 132 | 1,294 |
| Del-12 | 23 | 175 | 18 | 65 |
| Nz-1 | 409 | 9,869 | 113 | 1,348 |
| Sapporo-0 | 1 | 11,289 | 0 | 1,539 |
| Dra-2 | 408 | 9,893 | 82 | 1,365 |
| Belmonte-4-94 | 556 | 9,837 | 144 | 1,426 |
| Sha / Ara-1 | 244 | 11409 | 73 | 1,576 |
| Sij-2 | 1 | 11,287 | 0 | 1,560 |
| Wa-1 | 8 | 156 | 5 | 45 |
| **Col-0 (reference)** | **-** | **-** | **-** | **-** |
| AH-7 | 751 | 13,247 | 117 | 1,632 |
| Tibet-0 | 3,104 | 16,867 | 436 | 1,807 |
| Etna-2 | 2,234 | 14,988 | 341 | 1,681 |
| Meh-0 | 1,800 | 15,464 | 287 | 1,667 |
| Elk-1 | 1,164 | 13,188 | 196 | 1,527 |
| Ket-10 | 2,519 | 16,548 | 369 | 1,787 |
| Arb-0 | 310 | 12,494 | 233 | 1,549 |

**Supplementary Table 15** Summary of the structural variations (SVs) location detected in graph-based pan-genome constructed by 38 *Arabidopsis thaliana* ecotype genomes.

| **SV type** | **Gene Upstream 2k** | **Gene Downstream 2k** | **CDS** | **Intron** | **Total** |
| --- | --- | --- | --- | --- | --- |
| Biallelic/Insertion | 2,211 | 1,892 | 276 | 304 | 4,425 |
| Biallelic/Deletion | 1,497 | 1,440 | 447 | 473 | 3,316 |
| Biallelic/Divergent | 11,105 | 10,439 | 2,384 | 5,800 | 18,100 |
| Multiallelic | 7,672 | 6,513 | 1,584 | 2,188 | 12,423 |
| Total | 16,722 | 15,375 | 4,272 | 7,492 | 22,775 |

**Supplementary Table 16** Variance component analysis for 21 environment variables with SNPs and SVs.

| **Environment variables** | **SNPs variance component** | **SVs variance component** | **SNPs variance component in SNPs+SVs** | **SVs variance component in SNPs+SVs** | **SNPs+SVs variance component** |
| --- | --- | --- | --- | --- | --- |
| Annual mean UV-B | 0.983280008 | 0.919600986 | 0.776571091 | 0.205626815 | 0.982198 |
| BIO1 | 0.841911461 | 0.808743748 | 0.516222262 | 0.328525107 | 0.844747 |
| **BIO2** | **0.791628555** | **0.861090387** | **0.389532167** | **0.457891656** | **0.847424** |
| **BIO3** | **0.768939547** | **0.848347637** | **0.417842286** | **0.400443655** | **0.818286** |
| BIO4 | 0.850451653 | 0.851367639 | 0.478727059 | 0.383477954 | 0.862205 |
| BIO5 | 0.865701307 | 0.834369135 | 0.537720461 | 0.329956073 | 0.867677 |
| BIO6 | 0.851605548 | 0.822552958 | 0.489644926 | 0.360603401 | 0.850248 |
| BIO7 | 0.875670849 | 0.851659396 | 0.524294233 | 0.354573923 | 0.878868 |
| **BIO8** | **0.622742294** | **0.732501299** | **0.269548762** | **0.418664669** | **0.688213** |
| **BIO9** | **0.722796309** | **0.790020432** | **0.301731557** | **0.469425032** | **0.771157** |
| BIO10 | 0.827545912 | 0.797267763 | 0.529155895 | 0.304543218 | 0.833699 |
| BIO11 | 0.84066097 | 0.820566523 | 0.47573774 | 0.367745731 | 0.843483 |
| BIO12 | 0.649107526 | 0.627967814 | 0.36243647 | 0.279697984 | 0.642134 |
| BIO13 | 0.766733384 | 0.711248828 | 0.455153797 | 0.291190842 | 0.746345 |
| **BIO14** | **0.600333629** | **0.640721256** | **0.295063633** | **0.329421774** | **0.624485** |
| BIO15 | 0.78840305 | 0.776694942 | 0.425934552 | 0.375216677 | 0.801151 |
| BIO16 | 0.7062897 | 0.661598426 | 0.41257111 | 0.279275873 | 0.691847 |
| **BIO17** | **0.637626151** | **0.67173944** | **0.323223862** | **0.336873509** | **0.660097** |
| **BIO18** | **0.744802587** | **0.759518762** | **0.397119924** | **0.361627538** | **0.758747** |
| BIO19 | 0.674268752 | 0.650152089 | 0.368793738 | 0.30143824 | 0.670232 |
| Elevation | 0.973412105 | 0.953097941 | 0.71157784 | 0.282266977 | 0.993845 |

**Supplementary Table 17** Summary of the strong correlation SVs and related genes with different environment variables detected by SV-GWAS.

| **Environment variables** | **SV location** | **Related genes in 10k region** | **GWAS P-value** |
| --- | --- | --- | --- |
| BIO1 | Chr1:25118861 | AT1G62020, AT1G62030, AT1G62040, AT1G62045, AT1G62050, AT1G62060 | 9.07E-08 |
| BIO4 | Chr1:11993983 | AT1G33020, AT1G33030, AT1G33040, AT1G33050, AT1G33055, AT1G33060 | 1.82E-06 |
|  | Chr 2:17874513 | AT2G34010, AT2G34020, AT2G34030, AT2G34040, AT2G34050, AT2G34060, AT2G34070 | 8.04E-07 |
| BIO5 | Chr 1:13213890 | AT1G35660, AT1G35670, AT1G35680 | 2.36E-06 |
| BIO6 | Chr 1:25118861 | AT1G62020, AT1G62030, AT1G62040, AT1G62045, AT1G62050, AT1G62060 | 6.40E-10 |
|  | Chr 2:3693044 | AT2G06925 | 7.93E-07 |
| BIO7 | Chr 1:1199039 | AT1G04410, AT1G04420, AT1G04430, AT1G04440 | 1.41E-06 |
|  | Chr 1:11993983 | AT1G33020, AT1G33030, AT1G33040, AT1G33050, AT1G33055, AT1G33060 | 1.63E-06 |
|  | Chr 2:17874513 | AT2G34010, AT2G34020, AT2G34030, AT2G34040, AT2G34050, AT2G34060, AT2G34070 | 2.87E-06 |
| BIO9 | Chr 1:25241315 | AT1G62320, AT1G62330, AT1G62333, AT1G62340, AT1G62350, AT1G62360 | 7.73E-07 |
|  | Chr 5:12513269 | - | 2.46E-06 |
| BIO11 | Chr 1:25118861 | AT1G62020, AT1G62030, AT1G62040, AT1G62045, AT1G62050, AT1G62060 | 9.27E-10 |
|  | Chr 2:3693044 | AT2G06925, AT2G06960 | 1.74E-06 |
| Elevation | Chr 1:3141272 | AT1G09650, AT1G09660, AT1G09665, AT1G09680, AT1G09690, AT1G09700, AT1G09710, AT1G09720, AT1G09730, AT1G09740 | 9.70E-07 |

**Supplementary Table 18** Primers used for RT-qPCR.

| **Primer Name** | **Sequence (5’ to 3’)** |
| --- | --- |
| EIF4A-QPCR-F | CGTGACCCGTGATGATGAGA |
| EIF4A-QPCR-R | AGTACGGCAGAGCAAACACA |
| CCR1-QPCR-F | GGAACCGTACGGAATCCAGATGATC |
| CCR1-QPCR-R | CCTCGTAGTCCTGAAGATCTGCTTTG |
| KNAT3-QPCR-F | ATGGCGTTTCATCACAATCATCTCTCAC |
| KNAT3-QPCR-R | CTTCTTGGAAATGTTGTTGCTGTTGC |

**Figures**

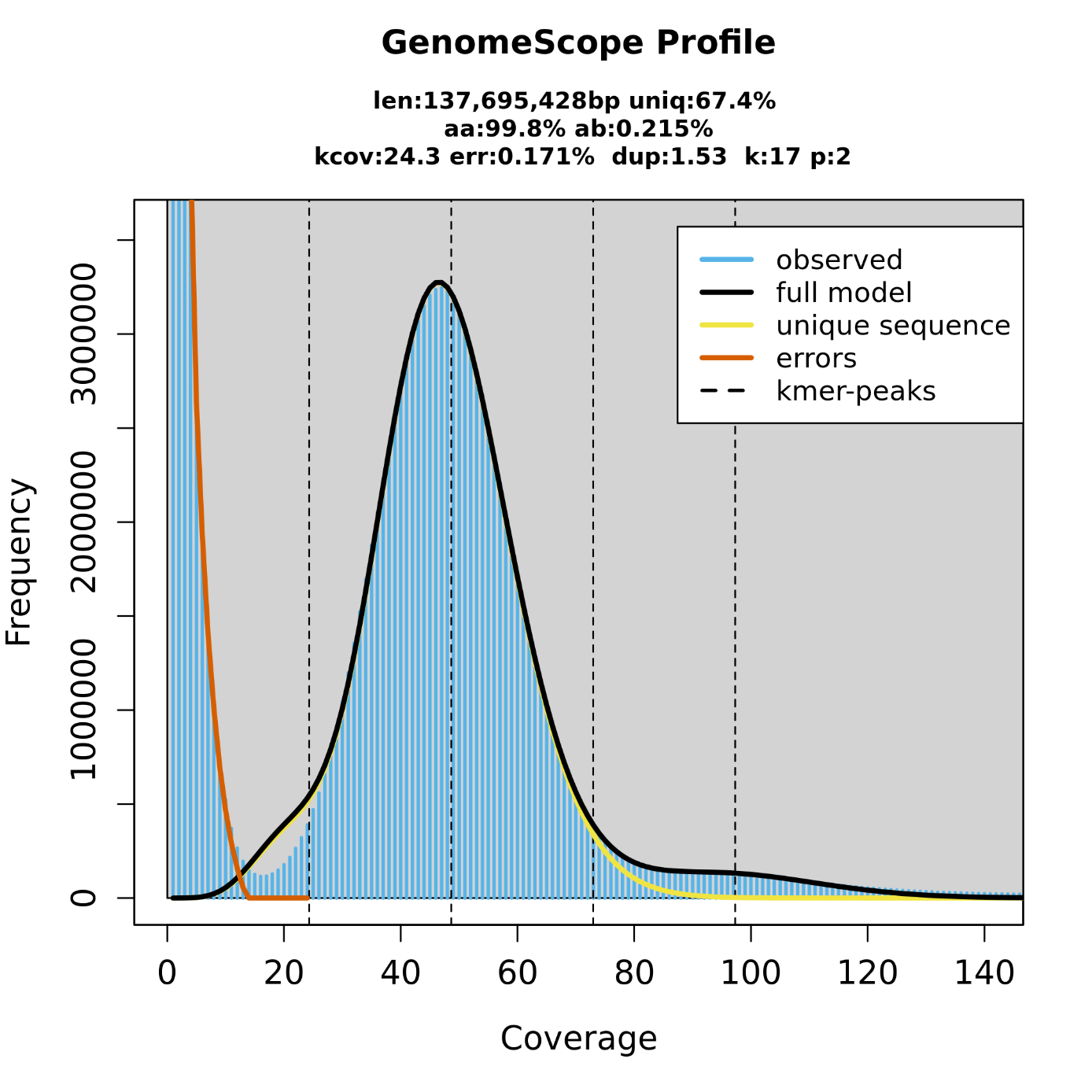

**Supplementary Figure 1** Estimation of Col-0 genome size by K-mer analysis. The figure shows the frequency of 17 k-mers, which are 17 bp sequences from clean reads of short-insert-size libraries. We identified 6,471,685,116 K-mers and the peak of K-mer depth is 47. Genome size can be estimated as (total K-mer number) / (the volume peak). The genome size of Col-0 was thus estimated as 137.70 Mb.

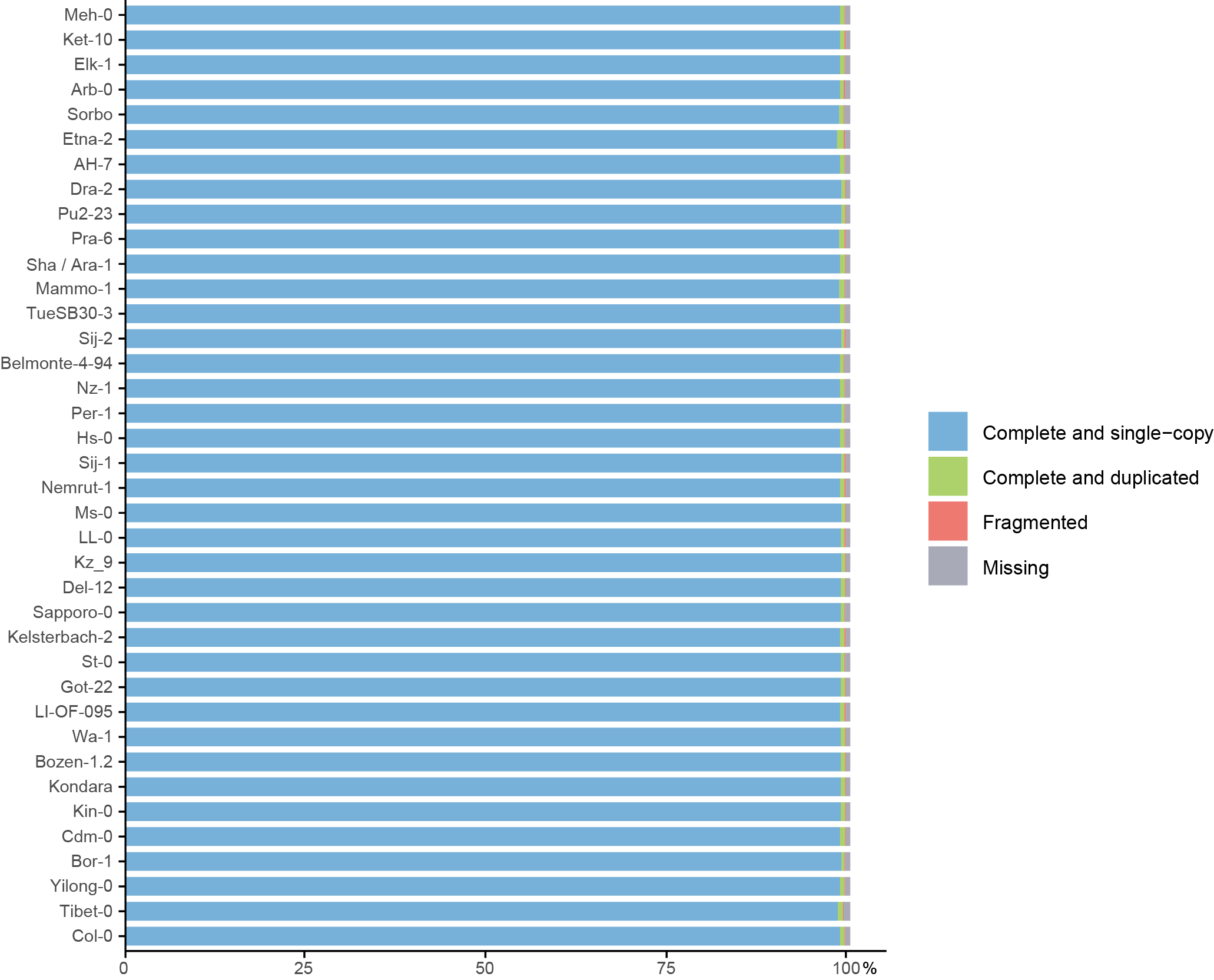

**Supplementary Figure 2** BUSCO assessment of the 38 *Arabidopsis thaliana* ecotypes genome assemblies.

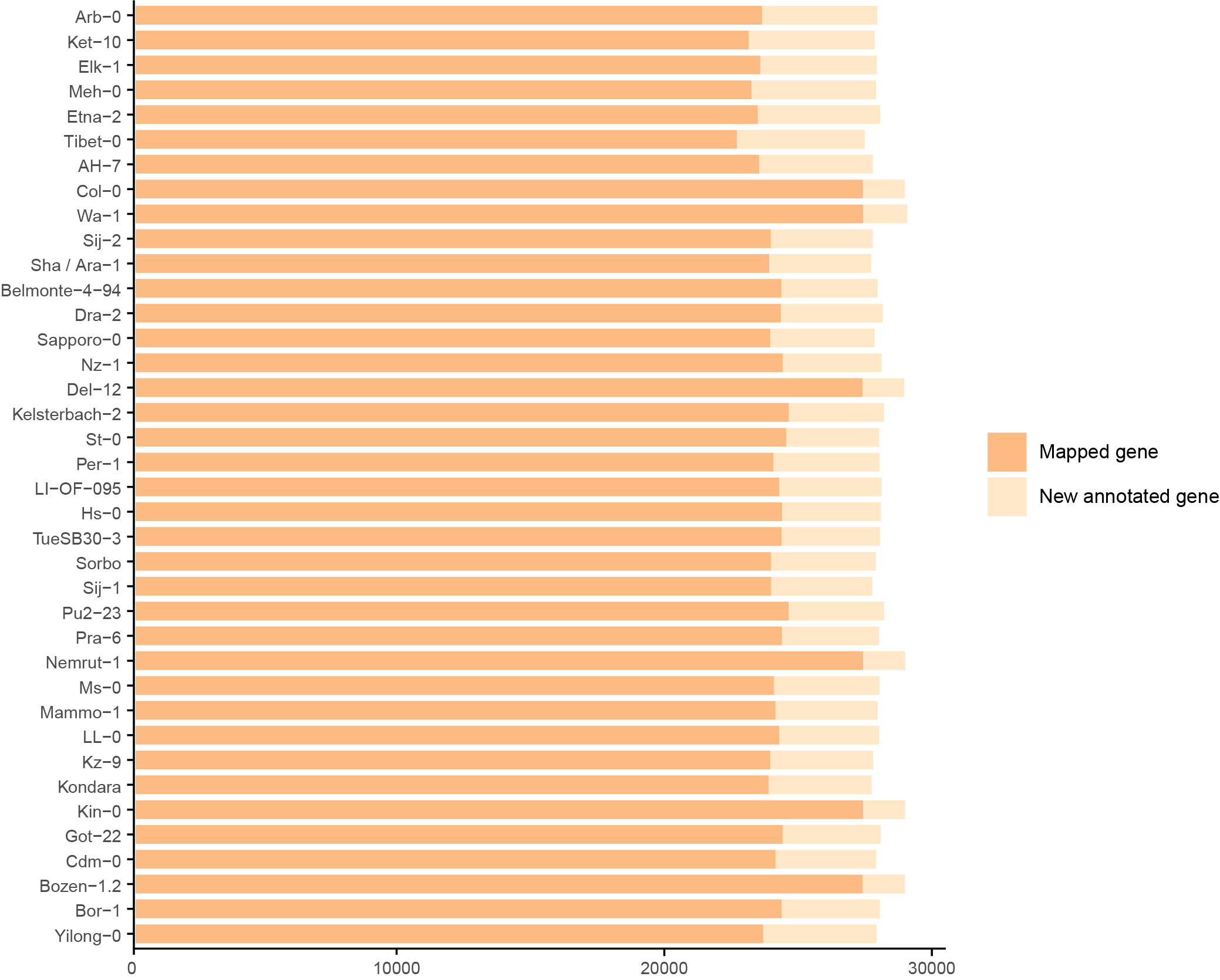

**Supplementary Figure 3** Summary of our 38 assemblies gene annotation compared with Araport11 version gene annotation of Col-0.

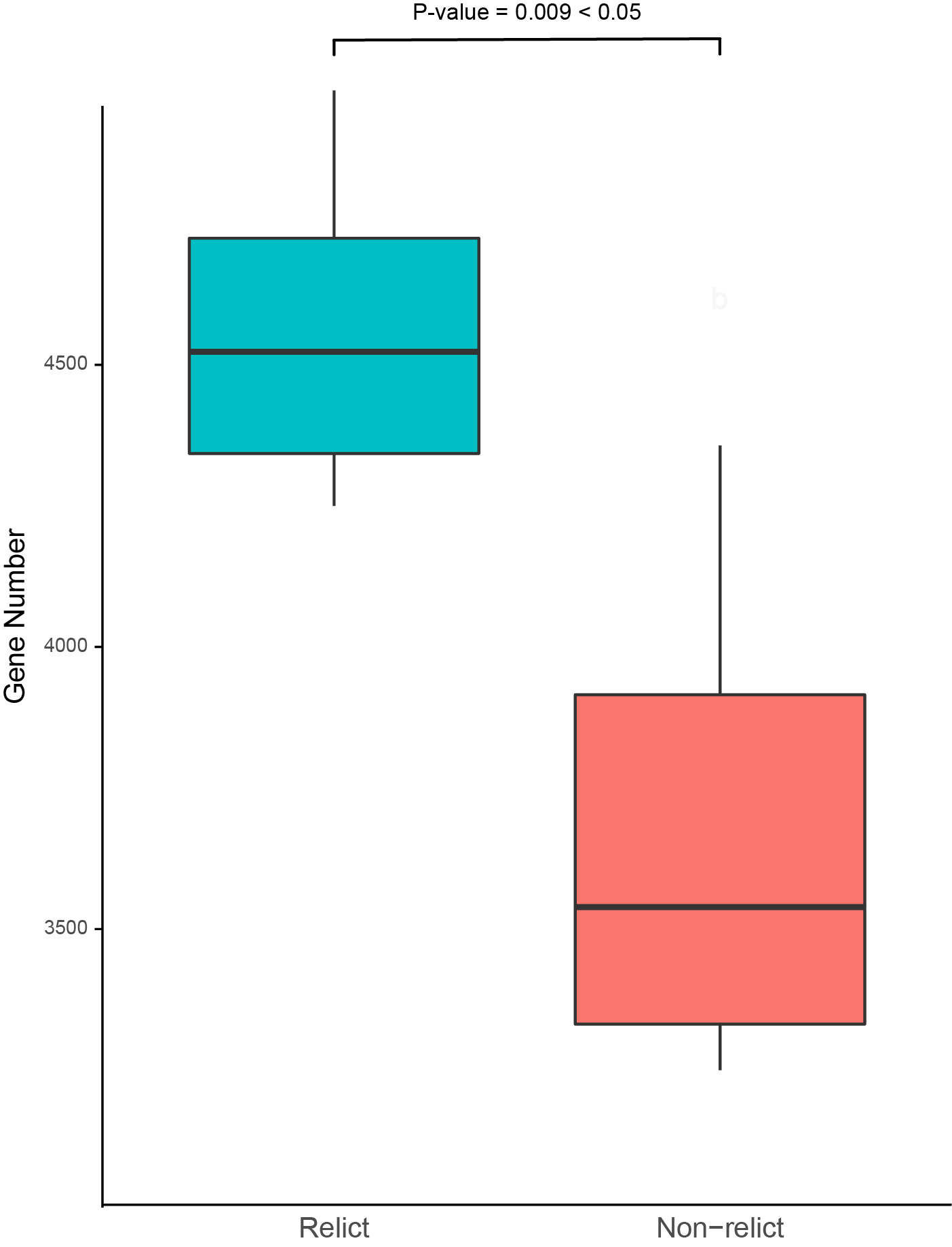

**Supplementary Figure 4** Comparison of genes different from Araport11 between relict ecotypes and non-relict ecotypes. Significance tested by Wilcoxon method with *p* = 0.00938915606307622 < 0.05. The middle line of the boxplot is the median, the lower and upper hinges correspond to the first and third quartiles, the upper whisker extends from the hinge to the largest value no further than 1.5 × IQR from the hinge (where IQR is the inter-quartile range) and the lower whisker extends from the hinge to the smallest value at most 1.5 × IQR of the hinge, the outliers are removed.

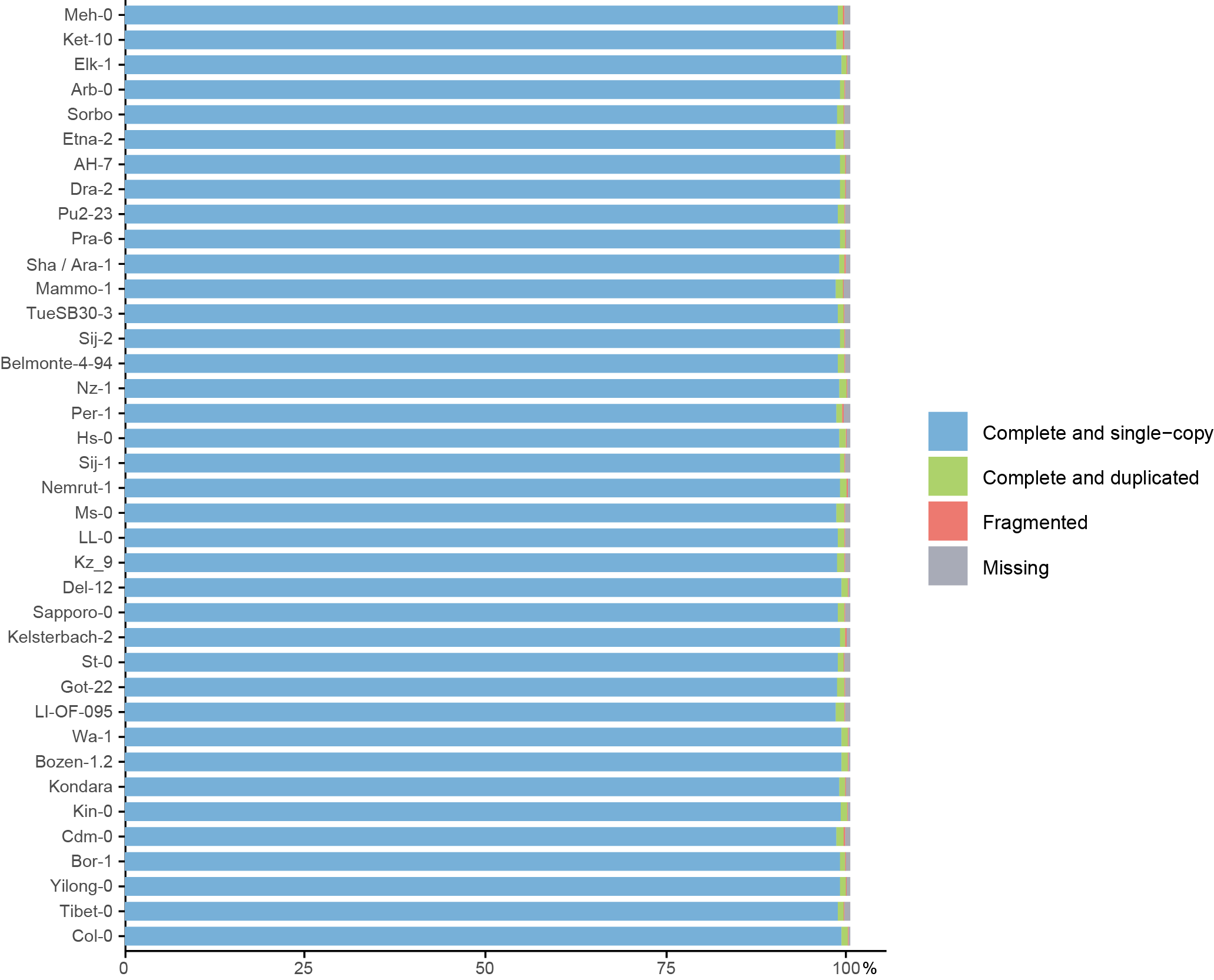

**Supplementary Figure 5** BUSCO assessment of the 38 *Arabidopsis thaliana* ecotypes genome assemblies’ gene annotations.

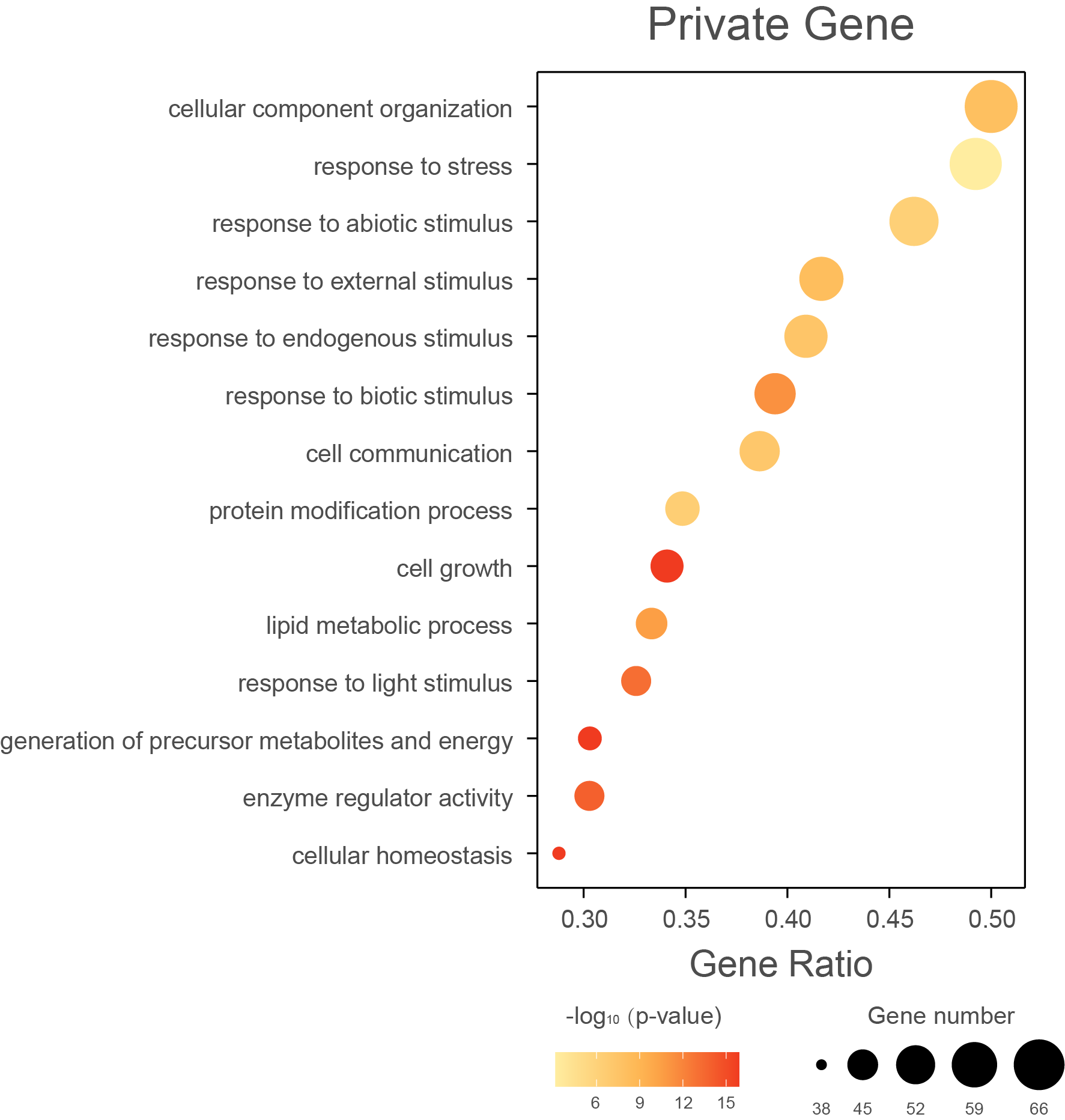

**Supplementary Figure 6** Bubble chart of GO enrichment analysis for private genes.

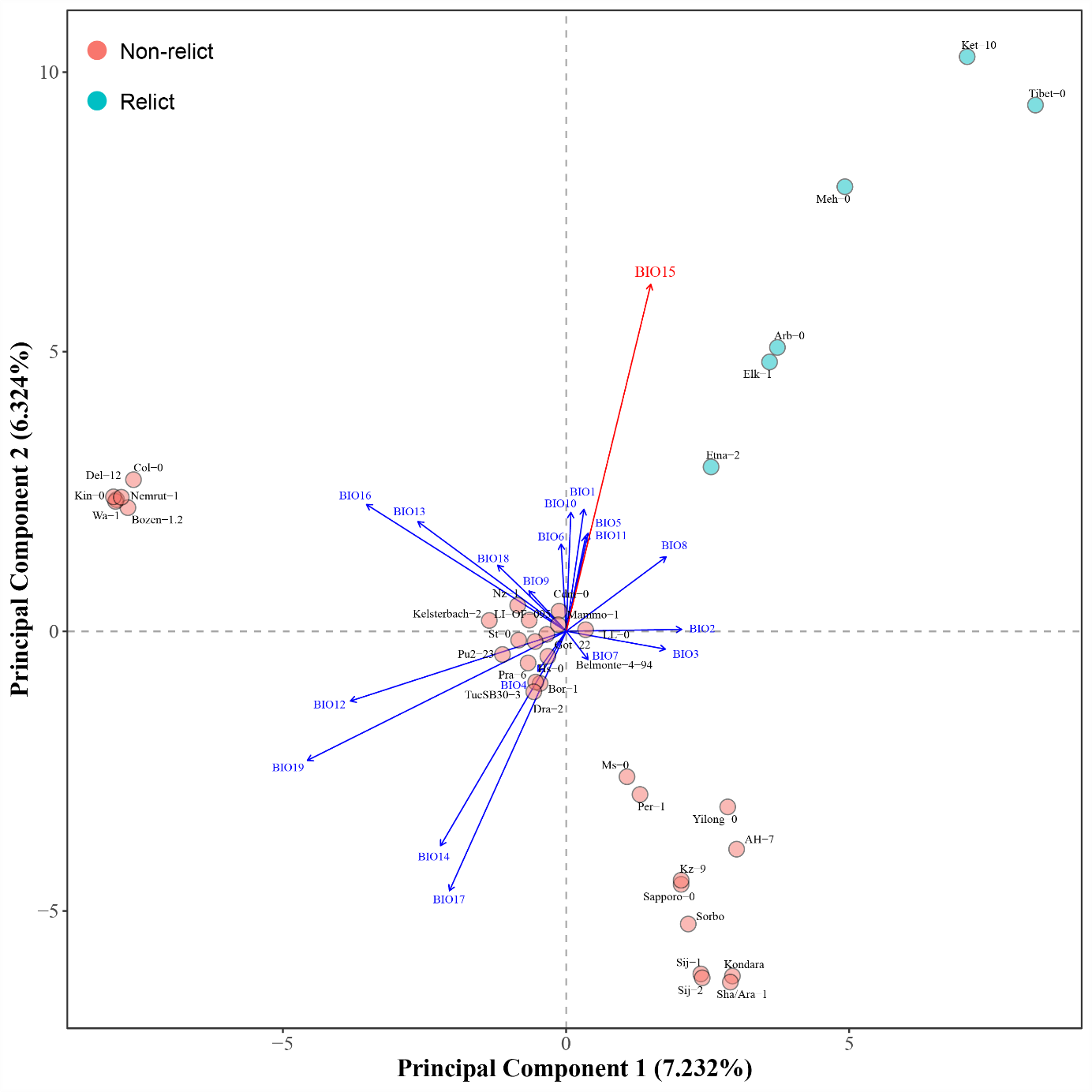

**Supplementary Figure 7** Principal component analysis (PCA) of 38 *Arabidopsis thaliana* ecotypes based on variable gene families and multiple regression of 19 BIOCLIM environmental variables on selected ordination axes. Blue circles displays relict ecotypes, while red circles displays non-relict ecotypes. Arrow in red shows significant assosiated environmental variables. Significance of the regression's coefficient of determination (r2) for each environmental variables was tested by 99,999 times Monte Carlo permutation test.

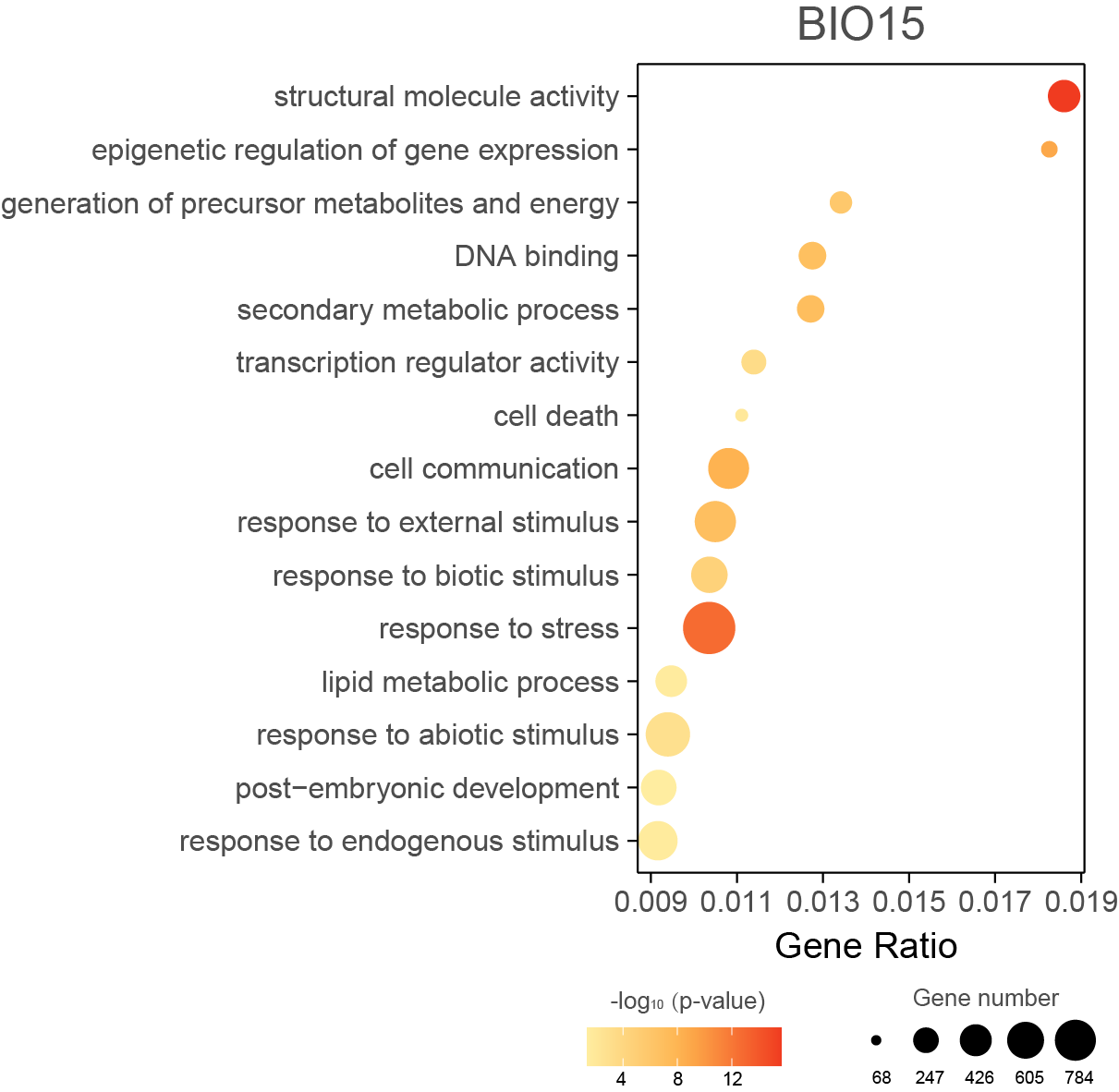

**Supplementary Figure 8** Bubble chart of GO enrichment analysis for 525 variable genes families significantly associated with BIO15.

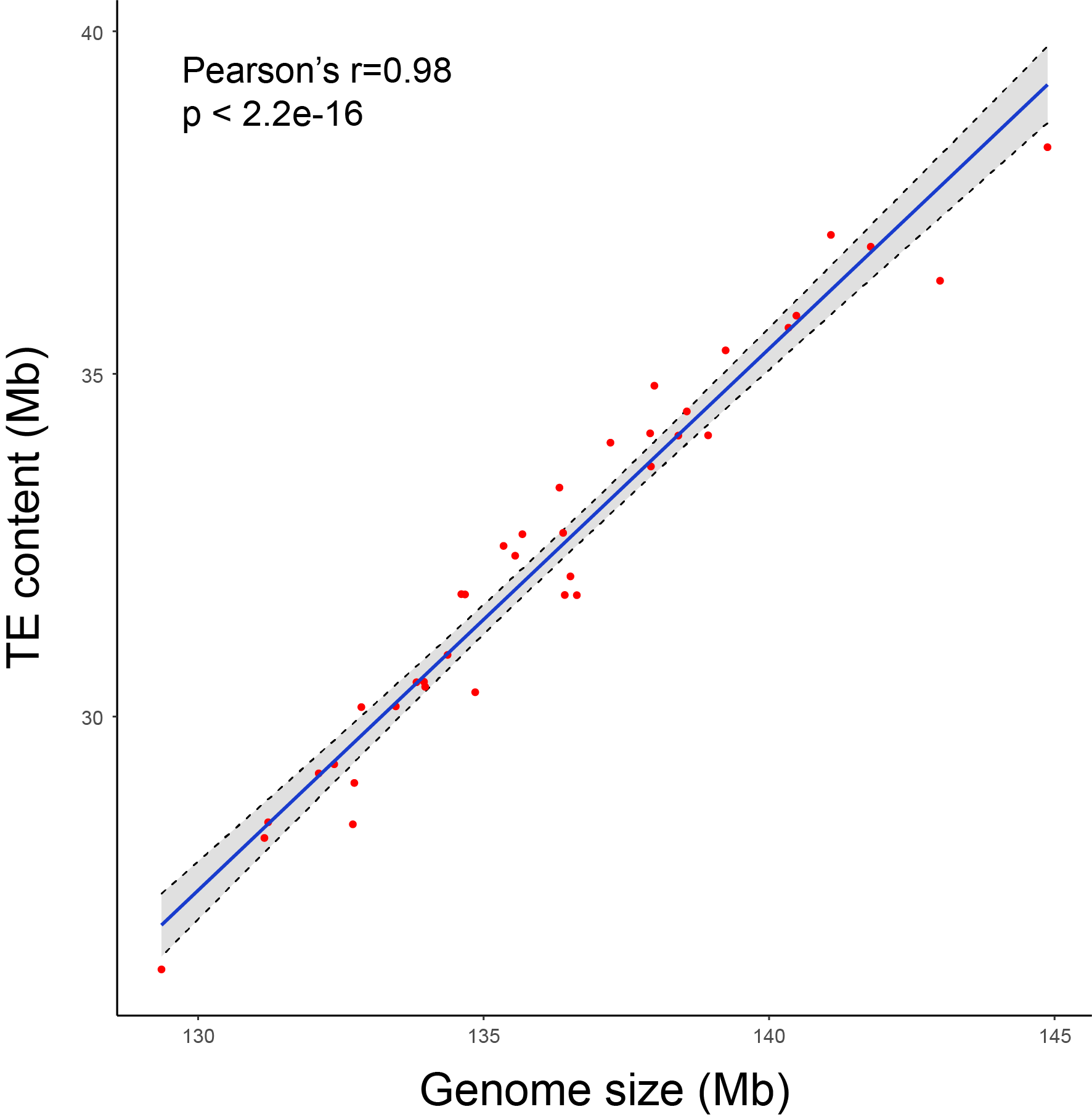

**Supplementary Figure 9** Pearson's correlation coefficient between TE content and genome size.

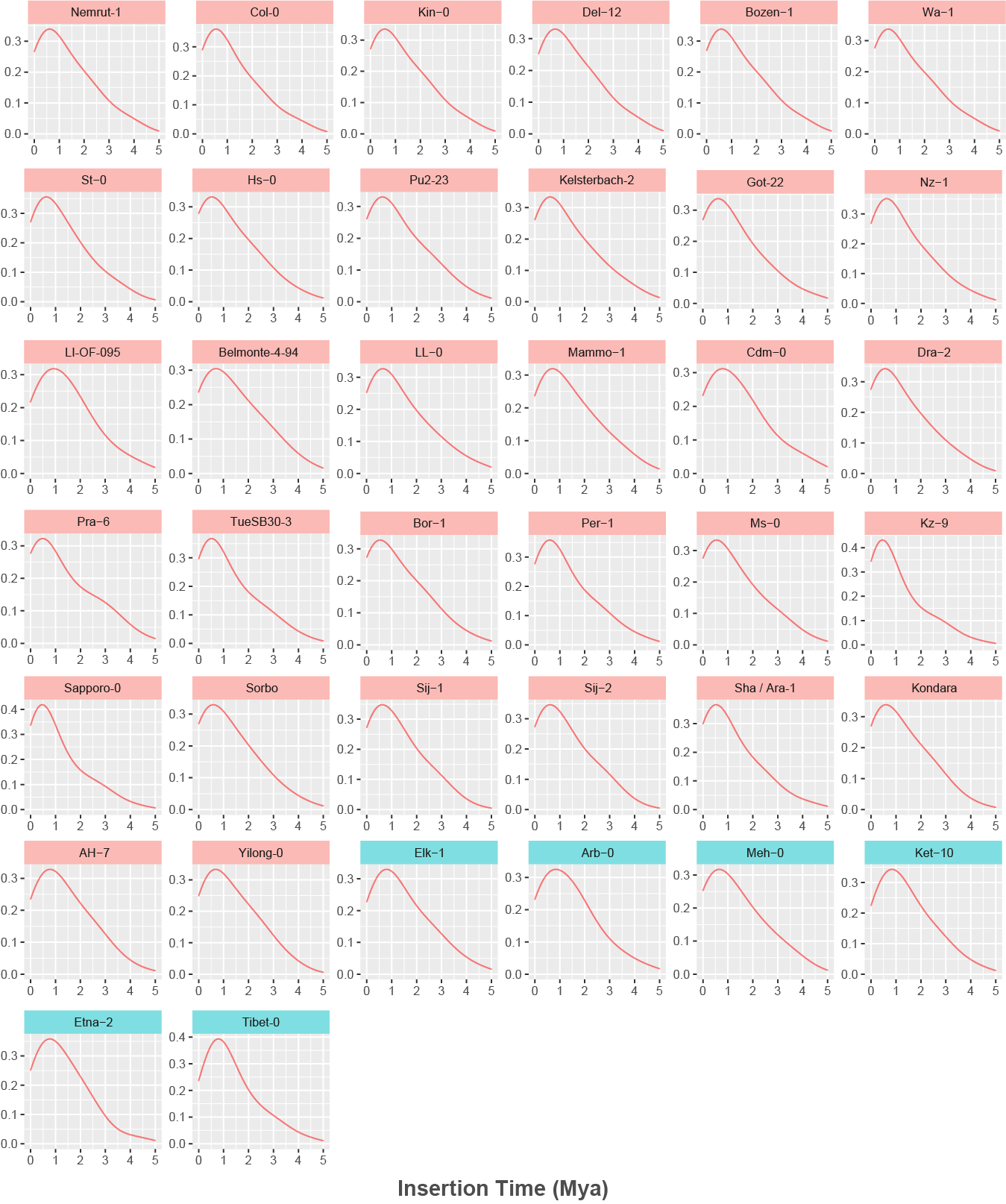

**Supplementary Figure 10** Intact LTR insertion time distribution of each accessions. The x-axis represents the insertion time from million years ago (Mya), y-axis represents the density of the insertion time. Blue rectangle displays relict ecotypes, while red rectangle displays non-relict ecotypes. The mutation rate used for time calculation was 7x10^-9^.

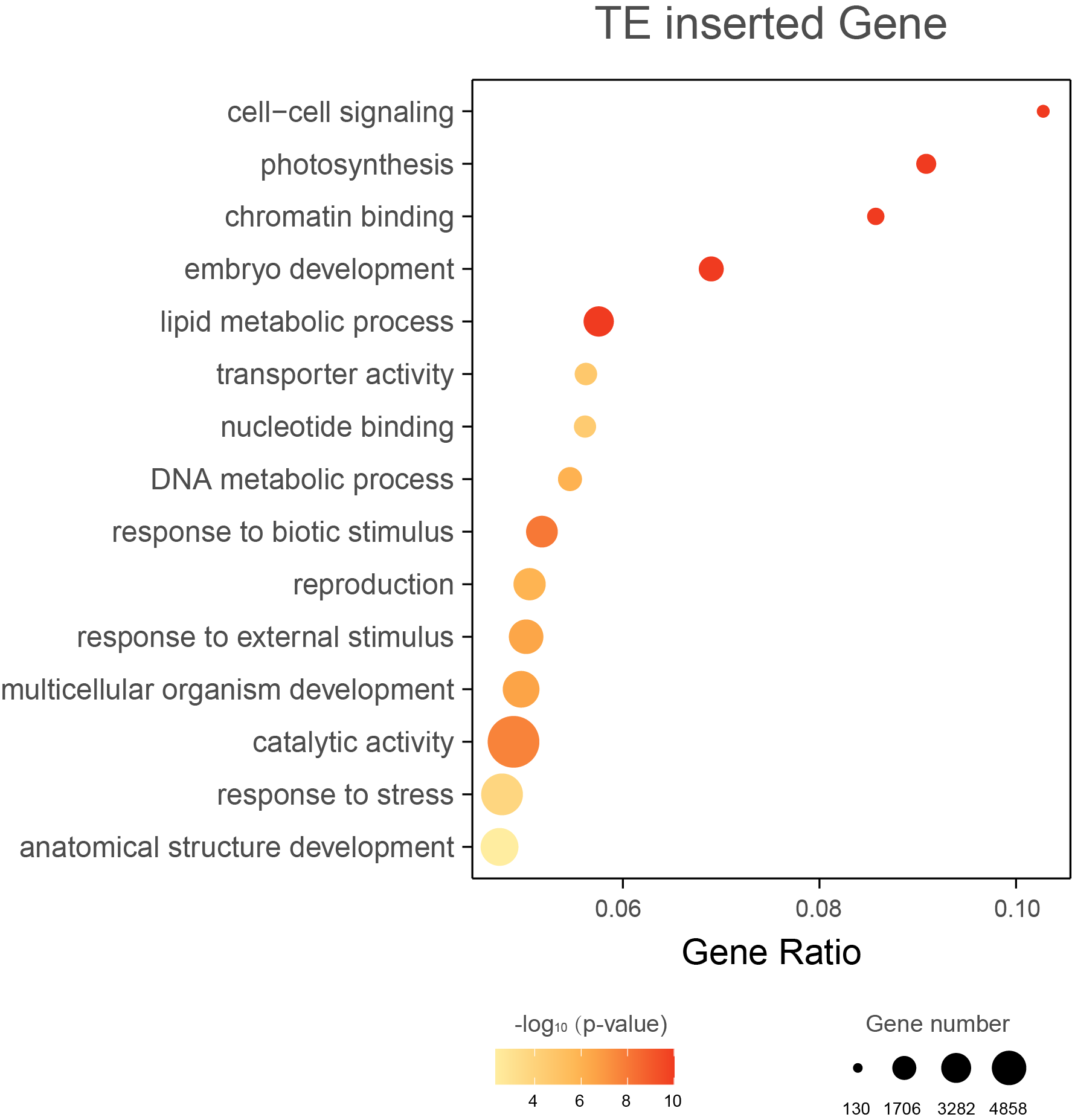

**Supplementary Figure 11** Bubble chart of GO enrichment analysis for TE inserted genes.

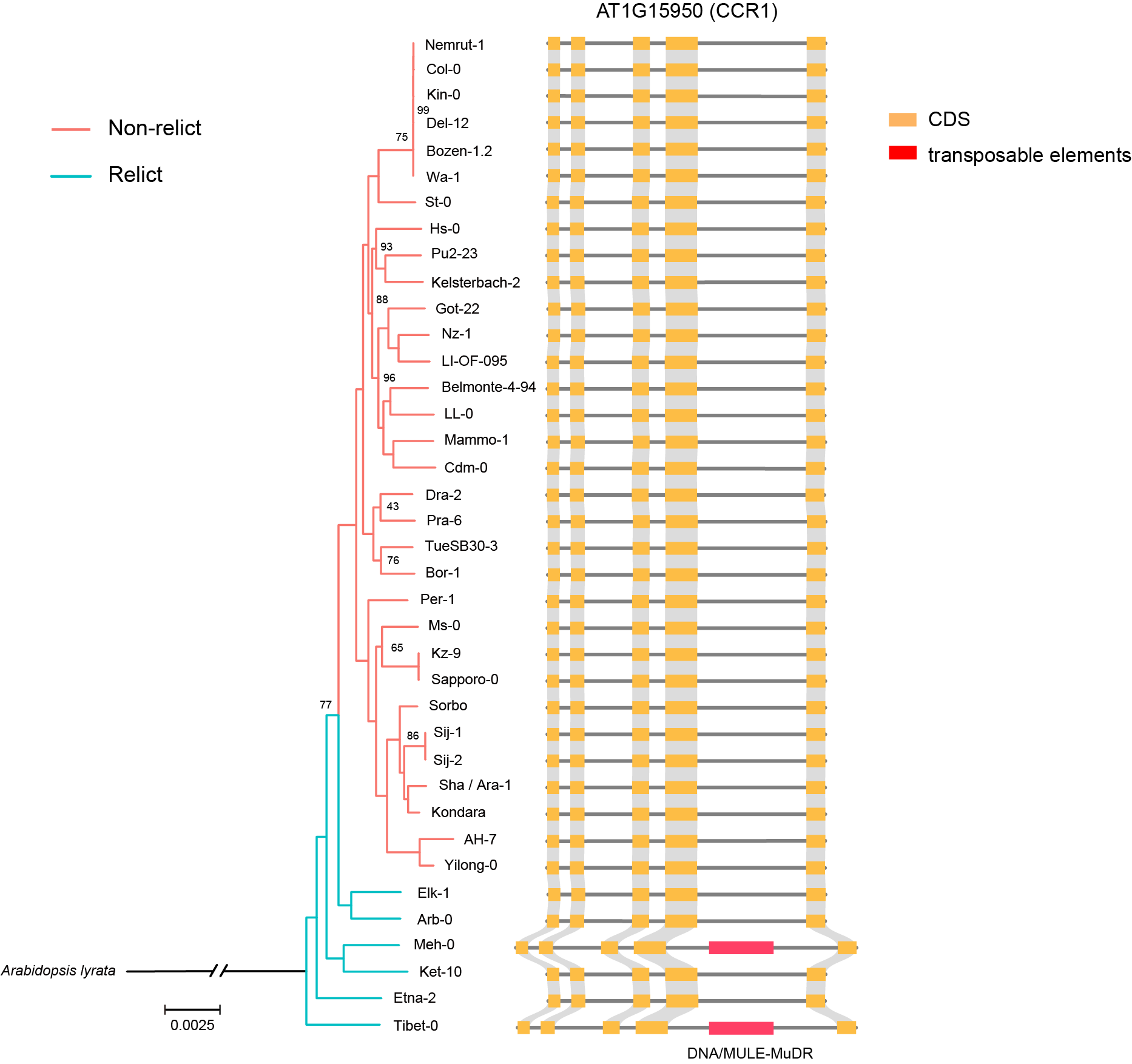

**Supplementary Figure 12** *AT1G15950 (CCR1)* gene structure in *Arabidopsis thaliana* ecotypes genomes. Only Tibet-0 and Meh-0 have a DNA transposon insertion in the fourth intron region.

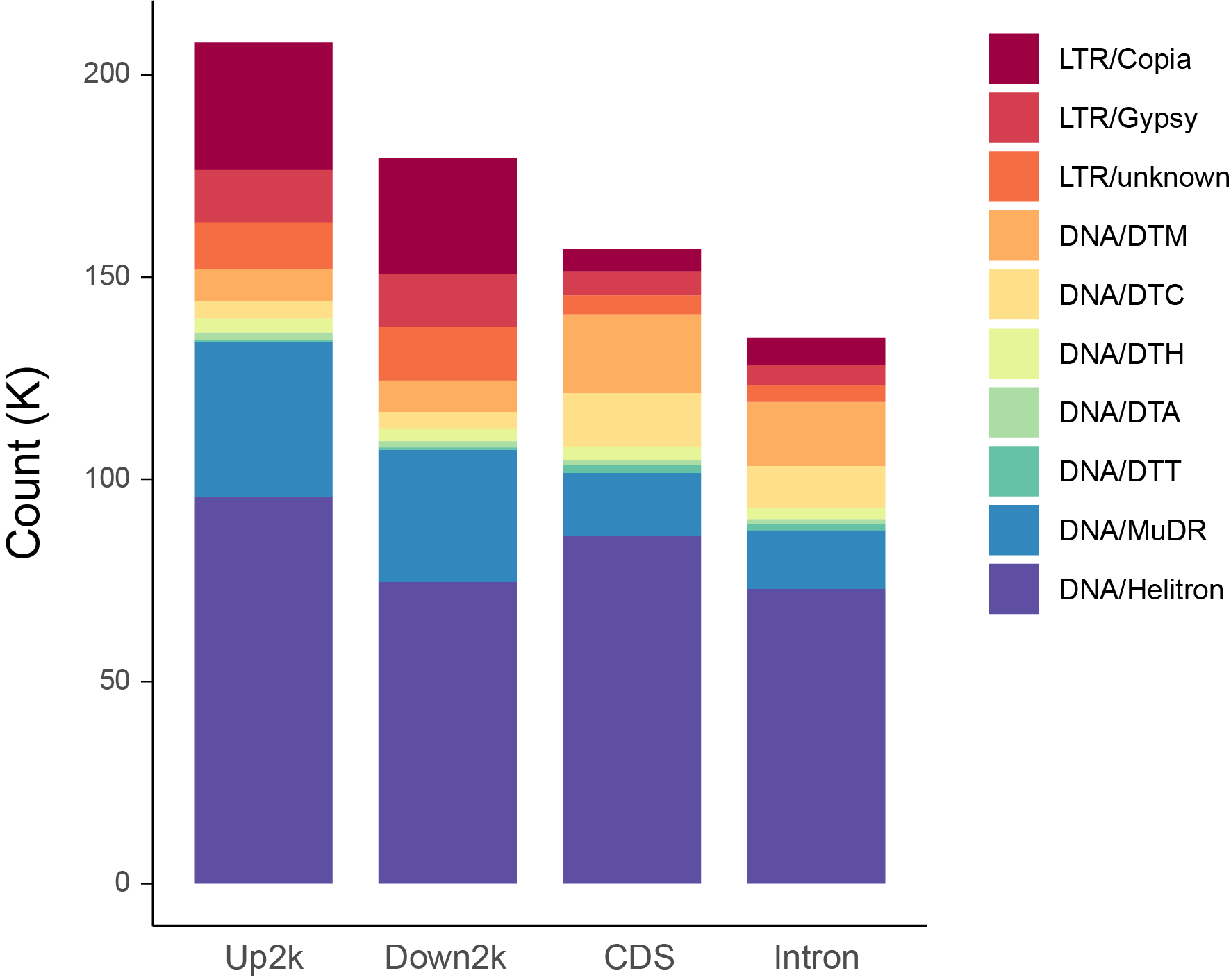

**Supplementary Figure 13** Composition of different TE types inserted around genes.

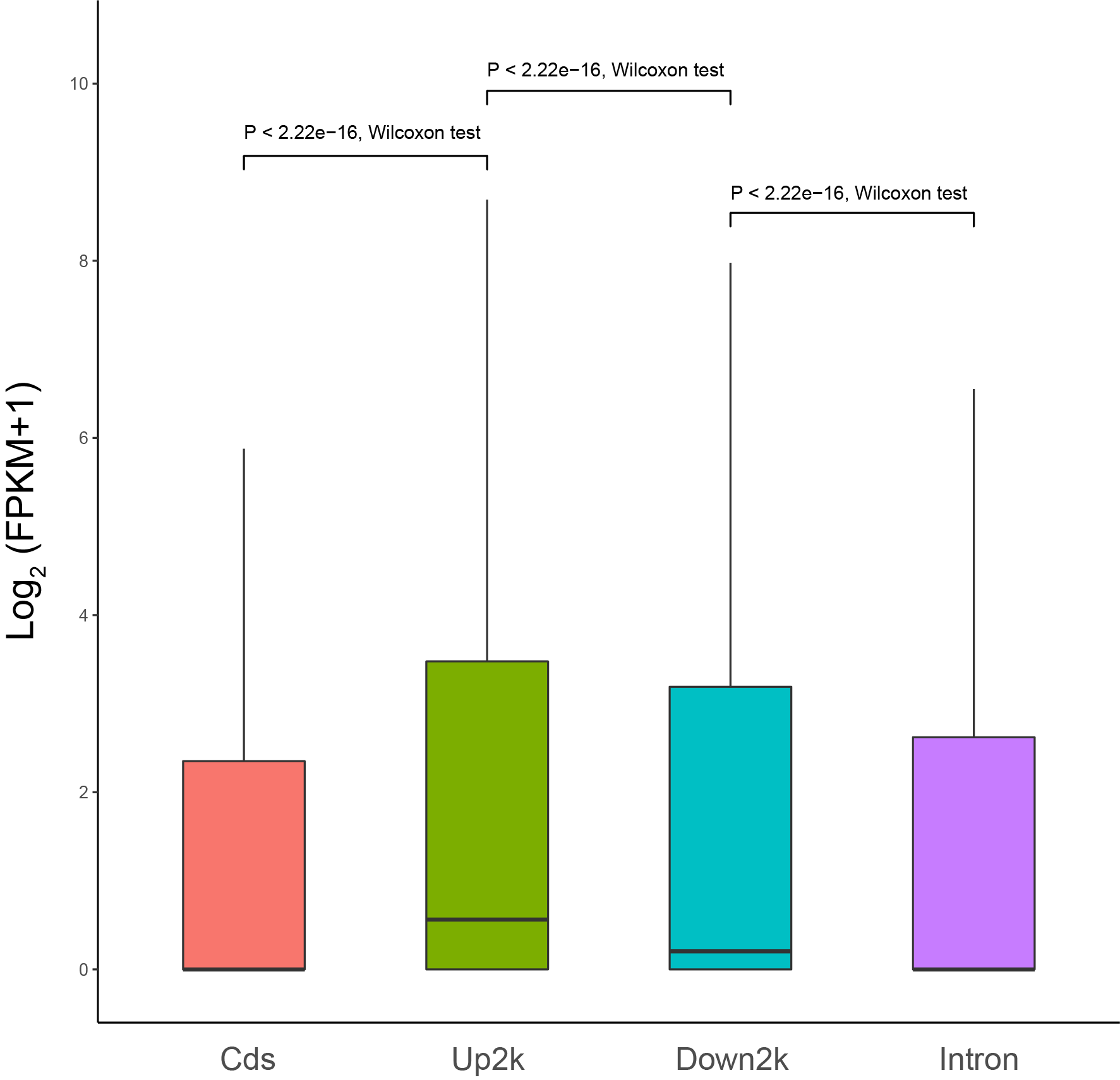

**Supplementary Figure 14** The expression level of genes with TE inserted in different regions. Significance tested by Wilcoxon method.

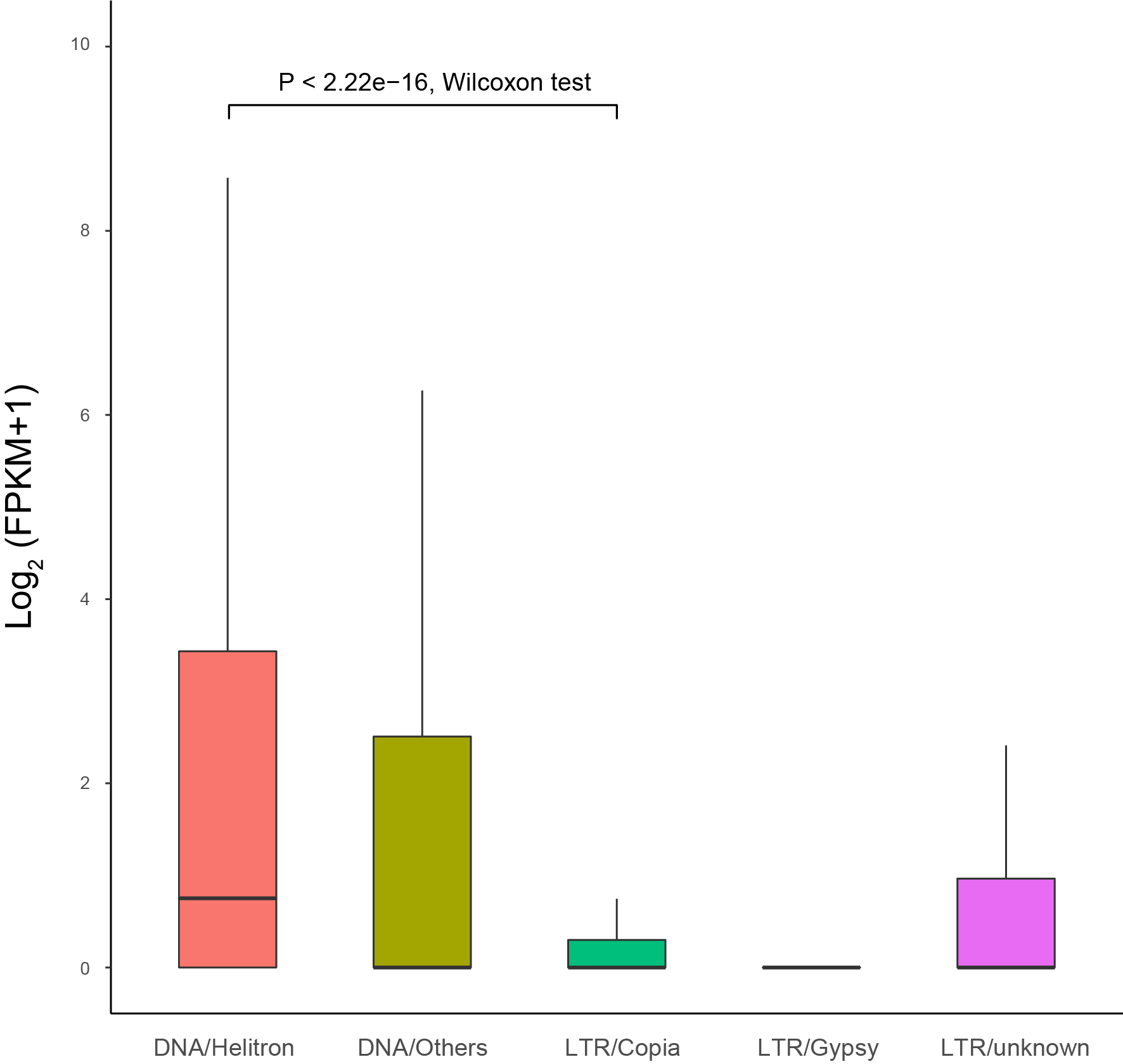

**Supplementary Figure 15** The expression level of genes with different TE types’ insertion. Significance tested by Wilcoxon method.

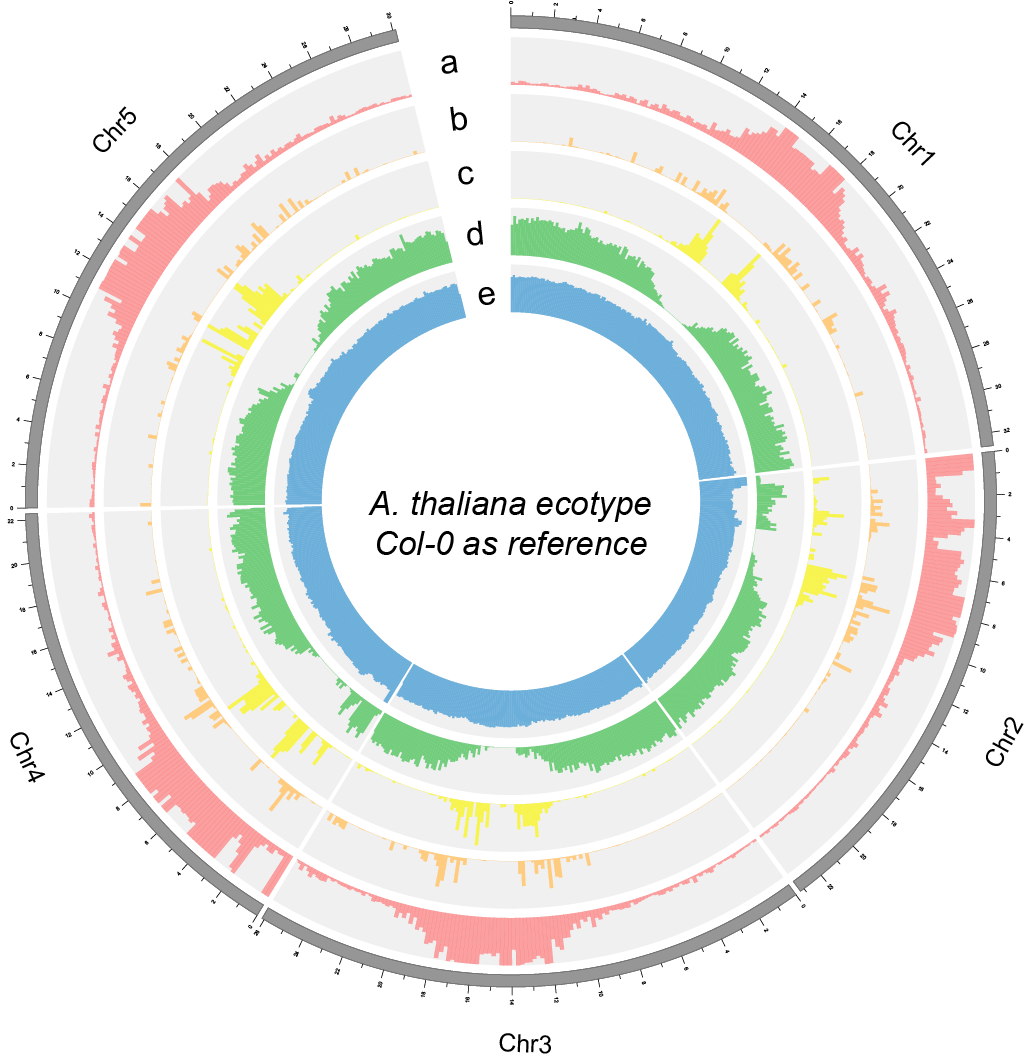

**Supplementary Figure 16** Circos plot of the reference genome Col-0. a. all TE distribution across the whole genome; b. LTR/Copia distribution across the whole genome; c. LTR/Gypsy distribution across the whole genome; d. gene density of the Col-0 genome; e. GC density of the Col-0 genome.

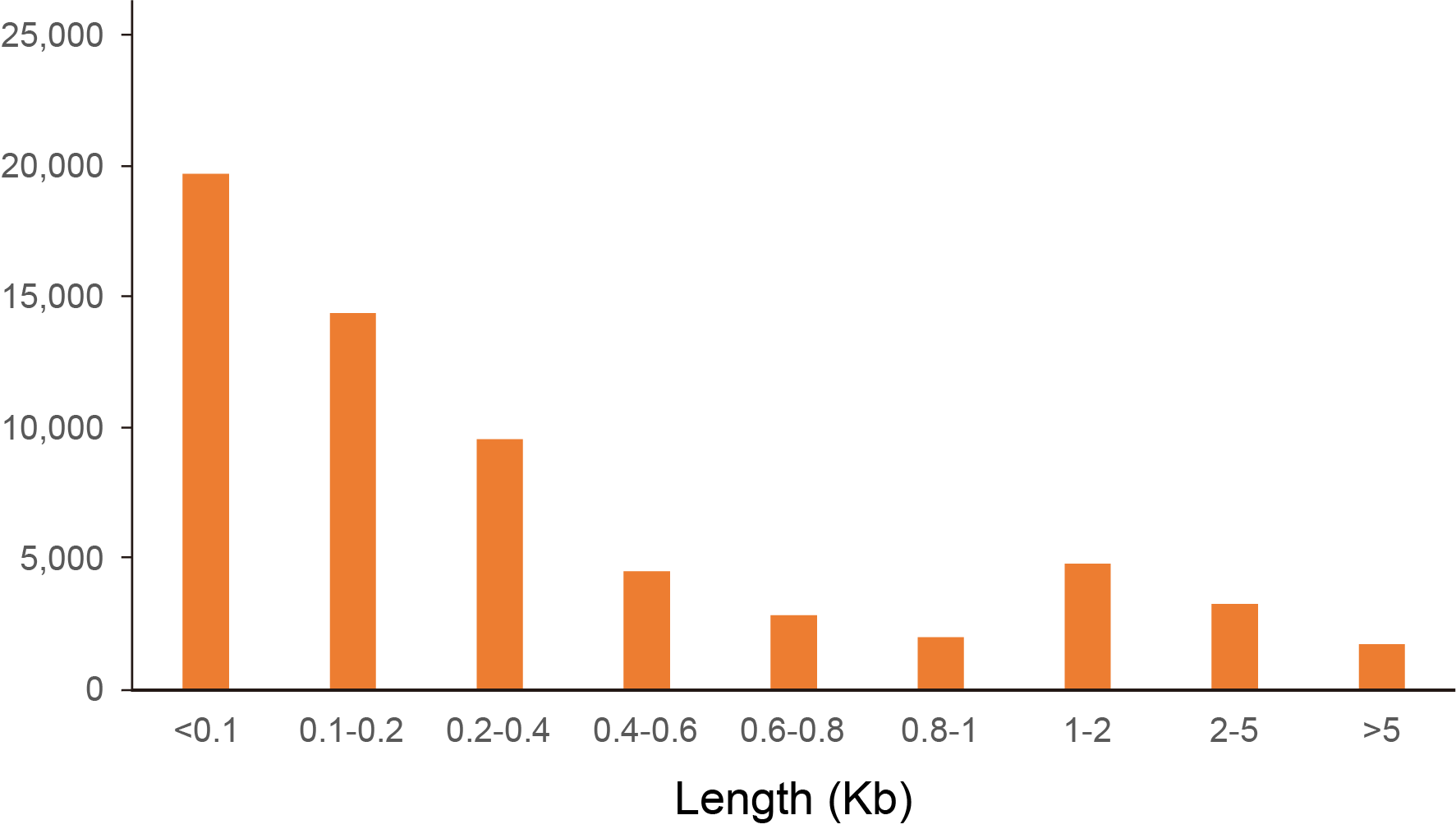

**Supplementary Figure 17** Distribution of the SVs length in graph-based pan-genome constructed by 38 *Arabidopsis thaliana* ecotype genomes.

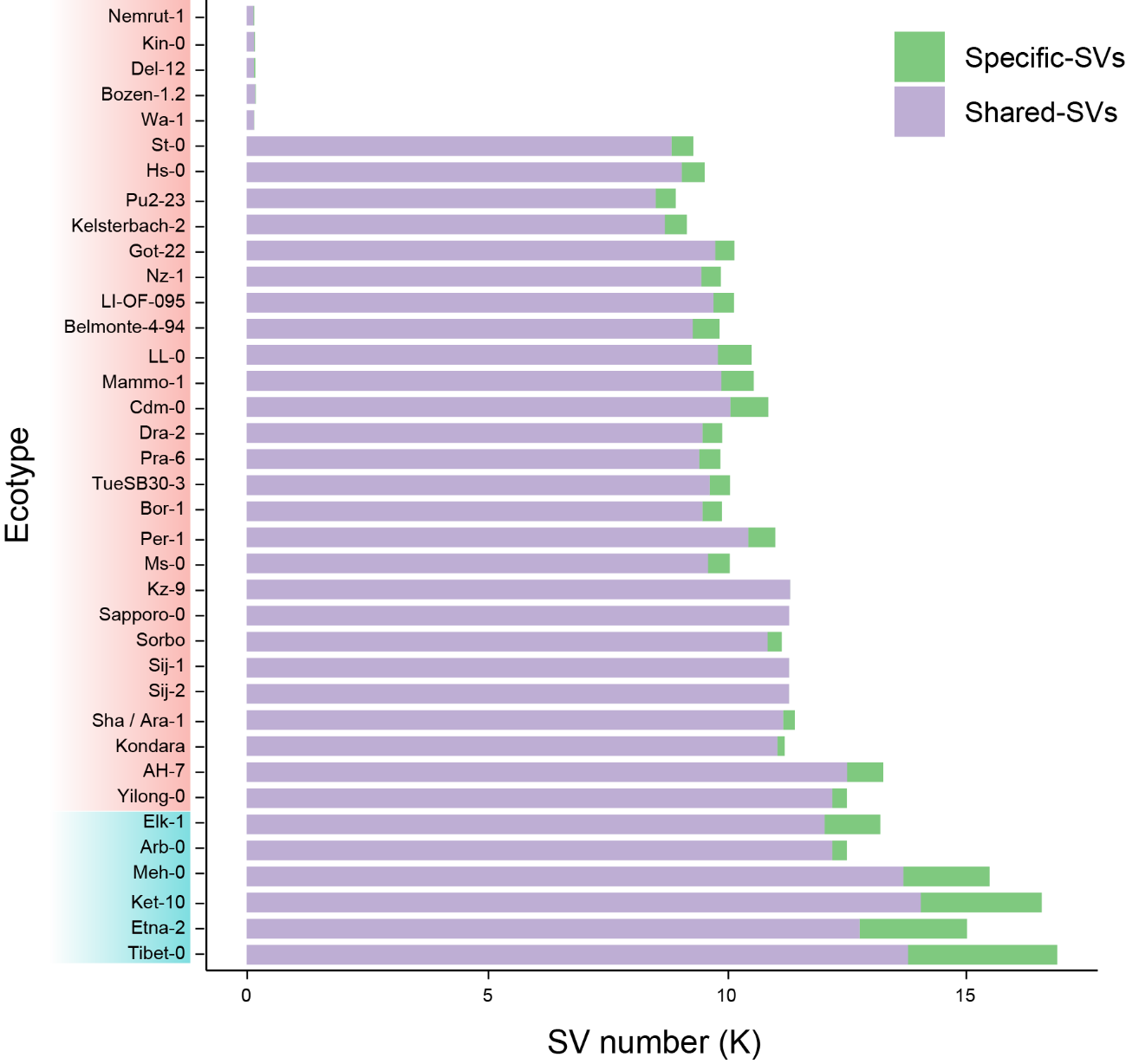

**Supplementary Figure 18** Shared-SVs and ecotype-specific SVs in 38 *A. thaliana* genomes. Blue rectangle displays relict ecotypes, while red rectangle displays non-relict ecotypes.

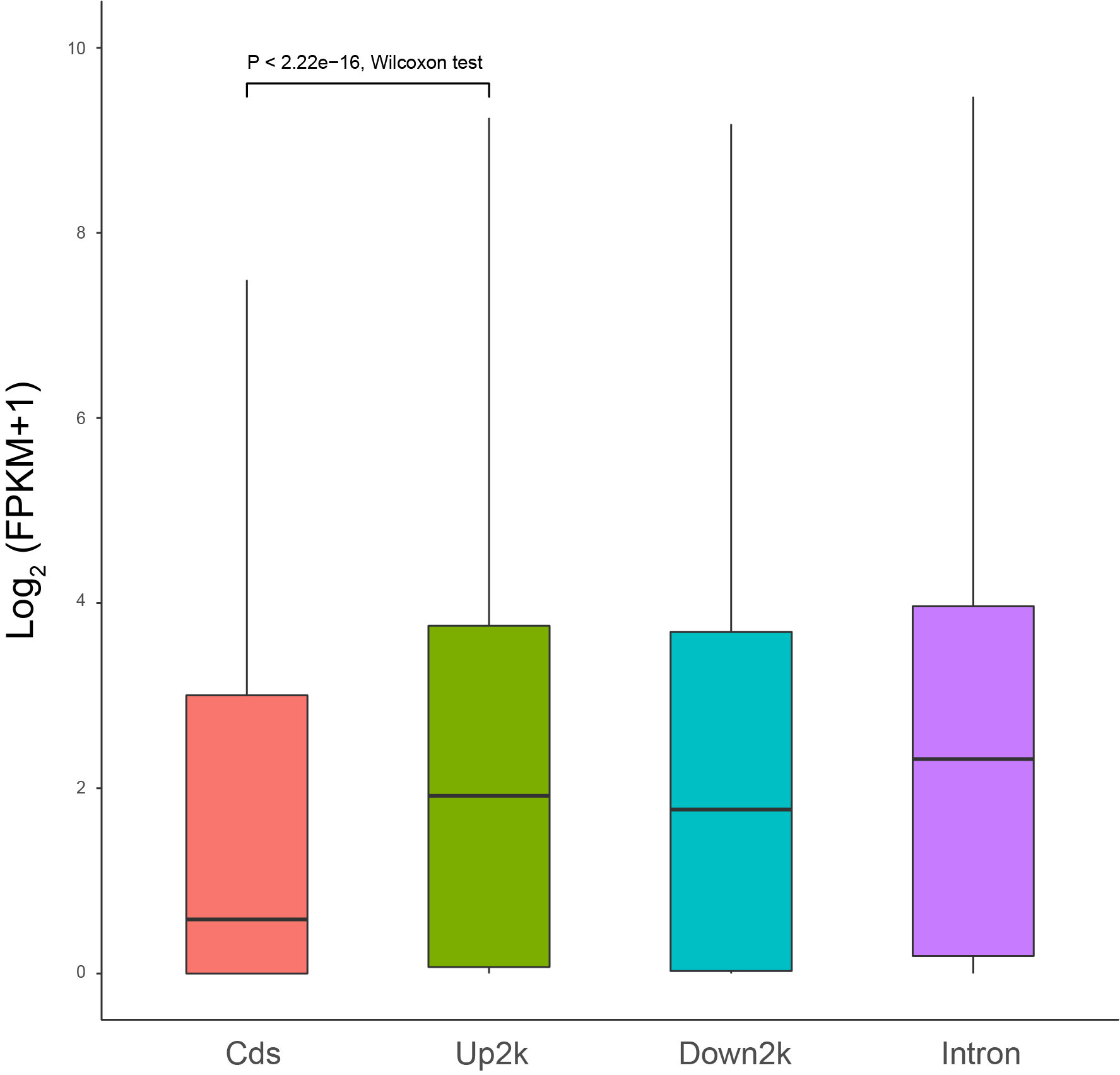

**Supplementary Figure 19** The expression level of genes with SV overlapped in different regions. Significance tested by Wilcoxon method.

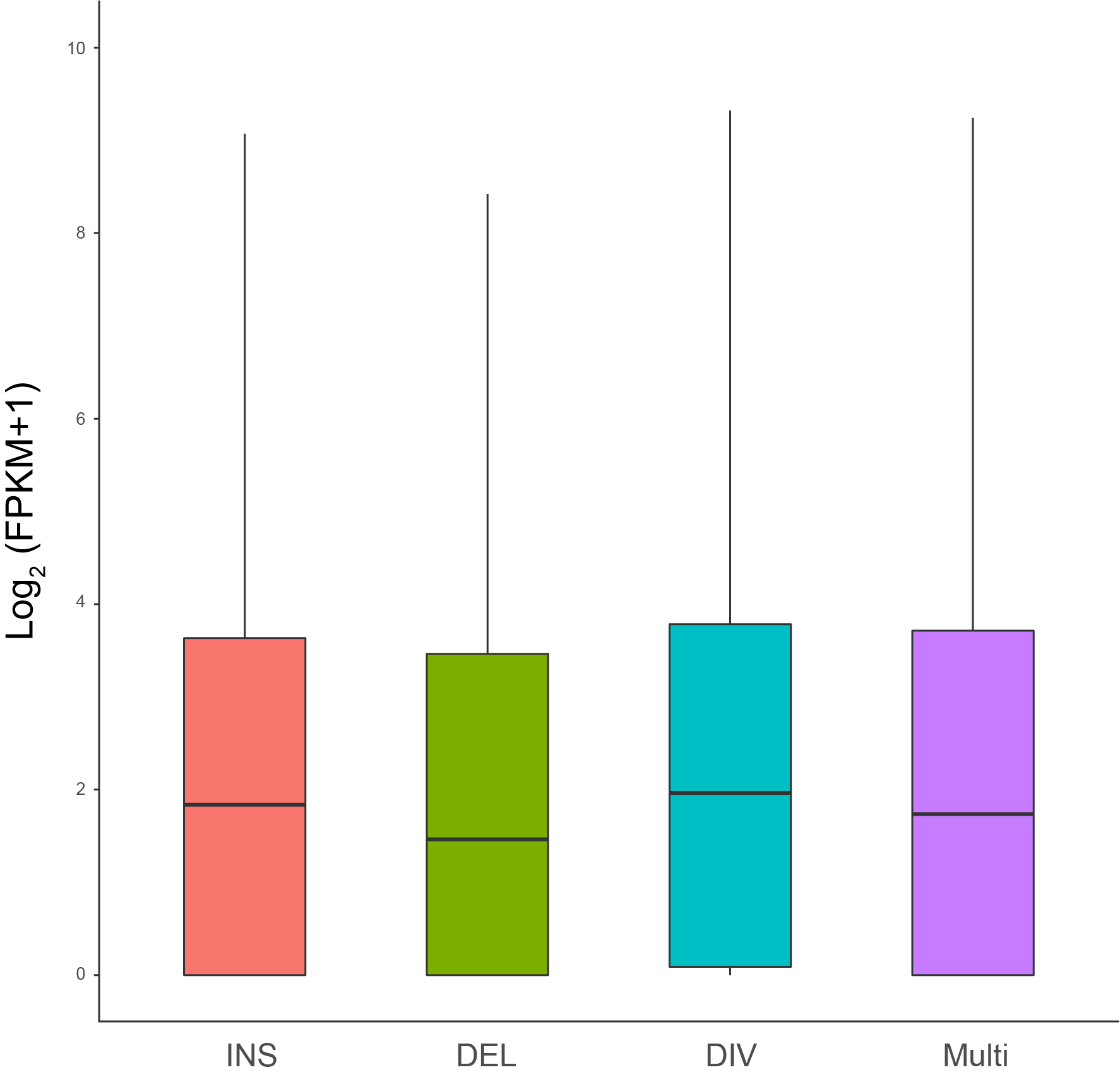

**Supplementary Figure 20** The expression level of genes overlapped with different SV types. Significance tested by Wilcoxon method.

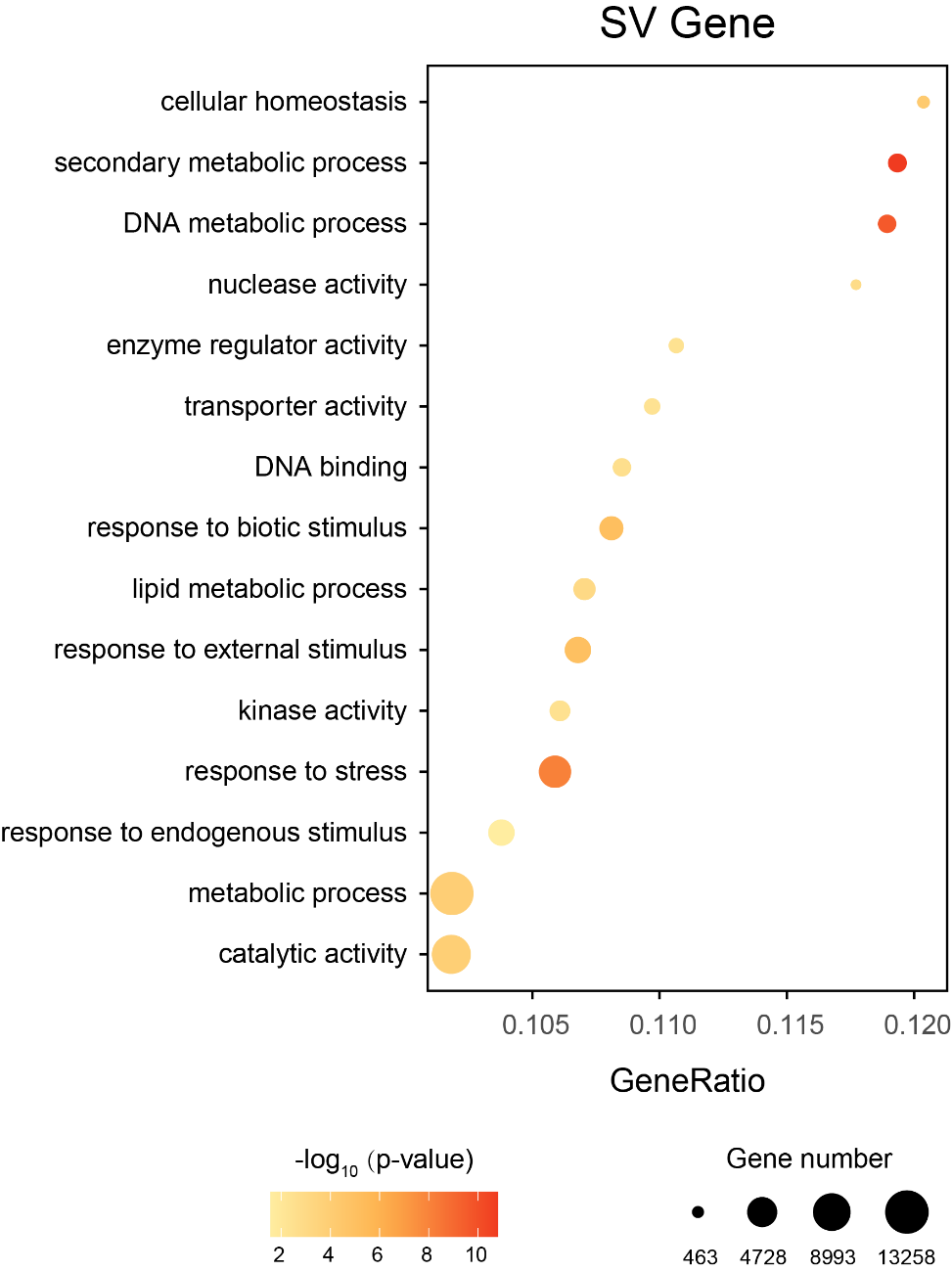

**Supplementary Figure 21** Bubble chart of GO enrichment analysis for SV overlapped genes.

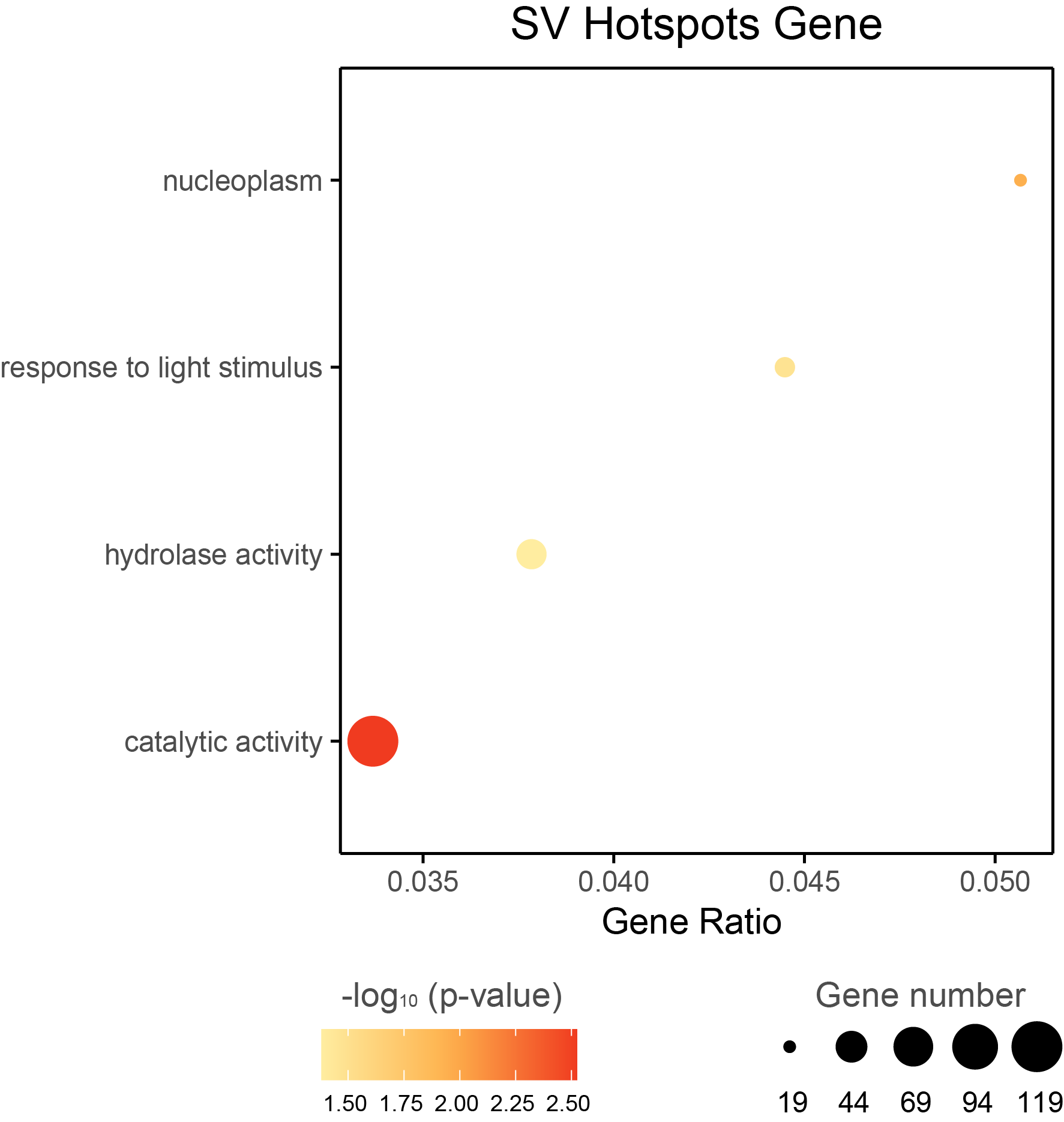

**Supplementary Figure 22** Bubble chart of GO enrichment analysis for genes in SV hotspot region.

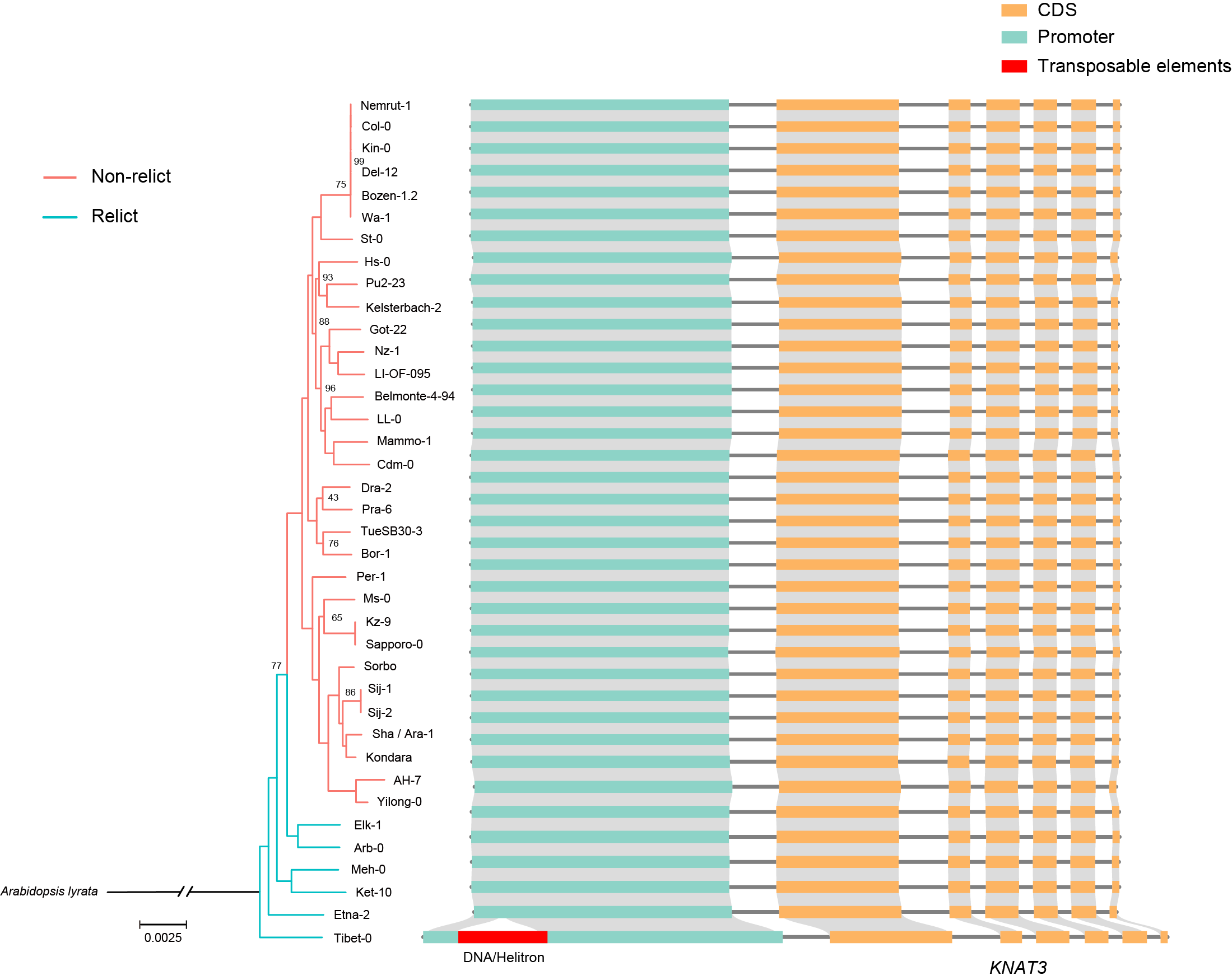

**Supplementary Figure 23** *AT5G25220 (KNAT3)* gene structure in *Arabidopsis thaliana* ecotypes genomes. Only Tibet-0 have a DNA transposon insertion in the promoter region.

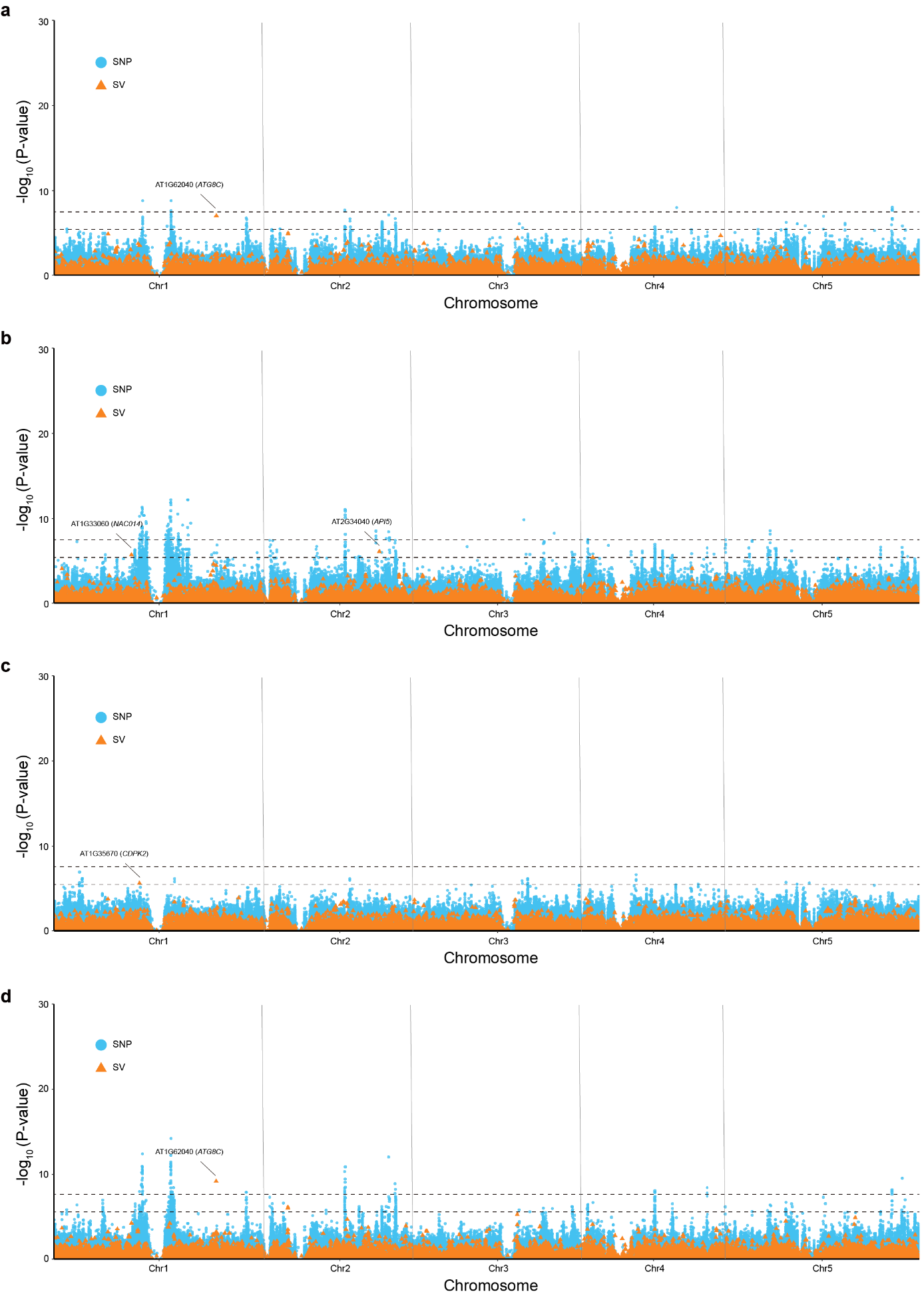

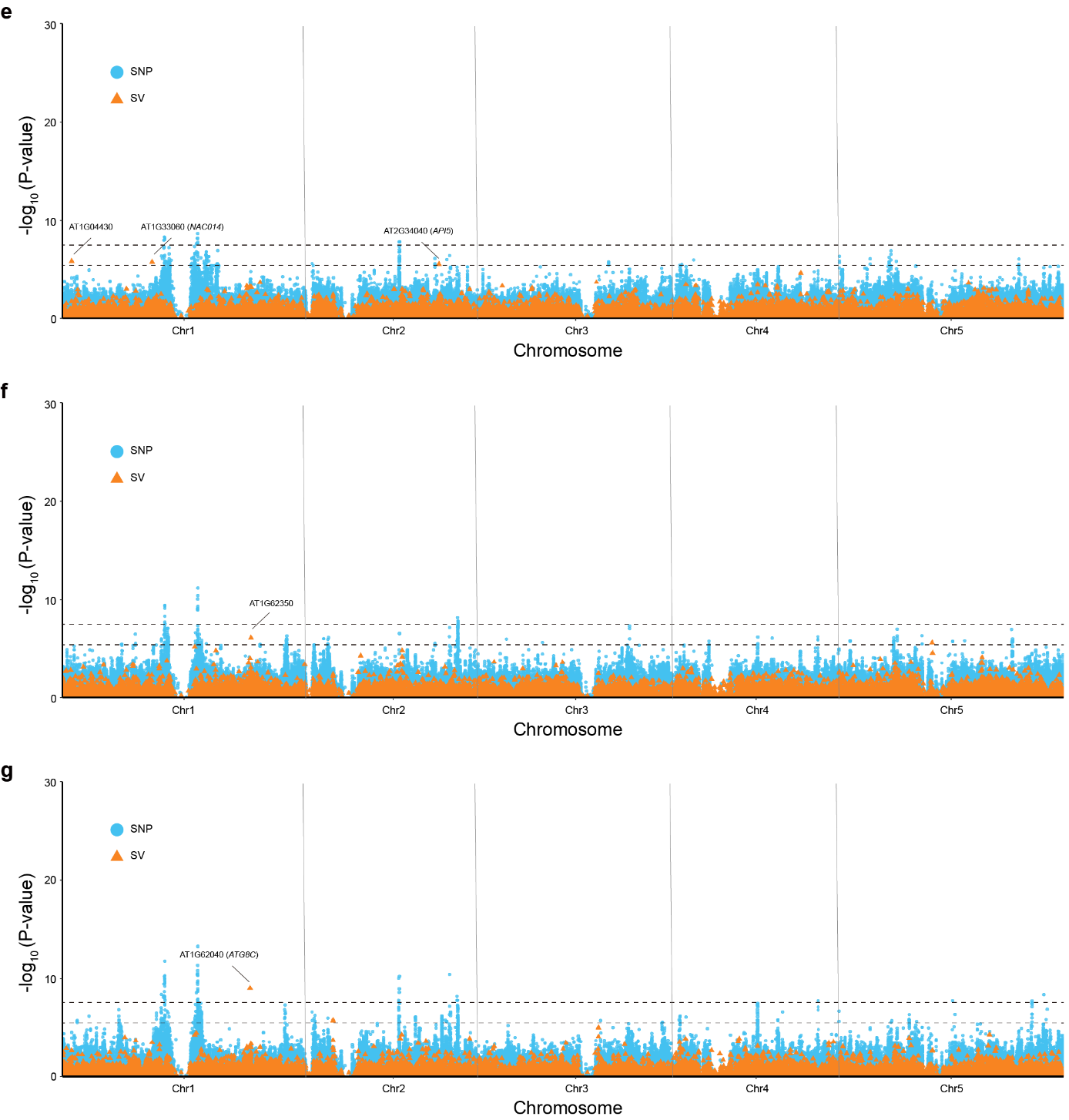

**Supplementary Figure 24** Manhattan plot of SV-GWAS (orange) and SNP-GWAS (blue) for environmental variables, including a. Bio1: annual mean temperature, b. Bio4: temperature seasonality (standard deviation ×100), c. Bio5: max temperature of warmest month, d. Bio 6: min temperature of coldest month, e. Bio7: temperature annual range (Bio5-Bio6), f. Bio9: mean temperature of driest quarter, g. Bio11: mean temperature of coldest quarter. The dashed black lines were genome wide significance threshold for SNP-GWAS (upper, 7.55) and SV-GWAS (lower, 5.47).
